# Cell therapy with IL-10-producing group 2 innate lymphoid cells suppresses Graft-versus-Host disease

**DOI:** 10.1101/2023.08.21.554158

**Authors:** Kyle T. Reid, Sarah J. Colpitts, Jessica A. Mathews, Abel Santos Carreira, Julia M. Murphy, Dorota T. Borovsky, Wenhui Cui, Tommy Alfaro Moya, Nadia Sachewsky, James An, Yubing Xia, Arthur Mortha, Jong Bok Lee, Li Zhang, Igor Novitzky-Basso, Jonas Mattsson, Sarah Q. Crome

## Abstract

IL-10 producing group 2 innate lymphoid cells (ILC2_10_) have immunoregulatory functions, and limit harmful immune responses across various tissues. Despite their crucial roles in maintaining immune homeostasis, the cell therapy potential of human ILC2_10_ has not been demonstrated, due to both limited numbers in human peripheral blood and lack of definitive markers for identification. Here, we isolate and expand circulating human ILC2_10_, and assess their cell therapy potential in a humanized model of Graft-versus-Host Disease (GVHD). Cell therapy with human ILC2_10_ decreased GVHD severity and prolonged survival of NOD-*scid* IL2Rγ^null^ (NSG) mice. Adoptive transfer of ILC2_10_ inhibited pathogenic T cell proliferation and intestinal infiltration, and suppressed CD4^+^ Th1 and CD8^+^ Tc1 cells in an IL-4 and IL-10 dependent manner. Critically, increased proportions of ILC2s did not correlate with higher rates of cancer relapse in HSCT recipients, and adoptive transfer of ILC2_10_ did not compromise graft-versus-leukemic (GVL) effects in a humanized model. Finally, we identify CD49d and CD86 as novel markers that discriminate ILC2_10_ from conventional ILC2s. Collectively, these findings demonstrate the potential of harnessing ILC2_10_ in cell therapies for GVHD and other immune-driven pathologies.

## Introduction

Group 2 innate lymphoid cells (ILC2s) have important roles in orchestrating immune responses and controlling tissue homeostasis. Initially described as a homogeneous population defined by expression of CD127, CD161 and CRTh2 in humans, recent findings have made it clear that ILC2s can differ in phenotype and function, both between and within tissues(1–4). For example, environmental signals can promote inflammatory ILC2s such as those that underly harmful immune responses in allergy and asthma, or instead promote immunoregulatory ILC2s that limit harmful inflammation in diverse contexts(5–8). While human ILC2 heterogeneity has been widely described, we are only beginning to appreciate how these distinct ILC2 populations interface with other immune cells, as well as how these different populations can be inhibited or harnessed in immunotherapy approaches.

IL-10-producing group 2 innate lymphoid cells (ILC2_10_) have been shown to inhibit immune responses in mouse models of inflammation and transplantation(9–14). For example, IL-2 administration induced IL-10 production in murine lung-resident ILC2s that have important functions in inhibiting and preventing lung inflammation (10), and intestinal mouse ILC2s were the primary producers of IL-10 at steady state and during disease(12). While there are few studies of human ILC2_10_, they are significantly reduced in individuals with allergies, but expand in response to allergy immunotherapy and are associated with treatment response(13), supporting human ILC2_10_ regulatory activity can be promoted with immunotherapy approaches to promote immune tolerance. The authors attempted to define unique ILC2_10_ markers, reporting that KLRG1 marked ILC2s that produced IL-10 along with other signature ILC2 cytokines(13). However, KLRG1 has also been widely reported on ILC2s with pro-inflammatory functions, limiting the use of KLRG1 as a unique marker for ILC2_10_(15–18). Therefore, identifying phenotypic features demarcating inflammatory and regulatory ILC2s is needed to enable comprehensive analysis of these functionally distinct ILC2 populations to health and disease, and to perform in depth studies of ILC2_10_ biology.

In addition to transplantation and autoimmunity, tolerogenic therapies are being widely pursued for the prevention and treatment of Graft-versus-Host Disease (GVHD) following allogeneic Hematopoietic Stem Cell Transplantation (HSCT). HSCT is a therapy commonly used to treat those with hematological malignancies, wherein transplanted donor stem cells develop into lymphocytes and attack cancerous cells, in what is termed the Graft-Versus-Leukemia(GVL) effect. While HSCT is often curative, up to 80% of patients develop acute GVHD characterized by donor T cells attacking healthy recipient tissues(19). Despite prophylactic use of post-transplant cyclophosphamide, acute GVHD remains the major cause of non-relapse associated morbidity and mortality, effecting 30-50% of HSCT recipients(20–24). Further, 30% of patients that develop GVHD fail to respond to corticosteroids or other broad immunosuppressive agents. Several regulatory cell therapy approaches have been explored in clinical trials, including adoptive transfer of regulatory T cells (Tregs). While evidence of Tregs limiting harmful allogeneic T cells have been reported, to date these cell therapies have only achieved limited clinical efficacy, and patients still require immunosuppression(25, 26). Therefore, there is an urgent need to develop more effective therapies for GVHD that do not impair GVL responses.

Prior studies reported ILCs are particularly sensitive to pre-HSCT conditioning therapies, where they are depleted from peripheral blood and tissues(27–29). Early Natural Killer (NK) cell reconstitution, in particular expansion of CD56^bright^ NK cells(30, 31), or the presence of activated ILC2s and group 3 ILCs(ILC3s) are associated with reduced GVHD incidence(32). Intriguingly, Bruce *et al.* showed that adoptive transfer of mouse ILC2s improved intestinal barrier function via ILC2-derived amphiregulin (AREG), and transferred ILC2s recruited and activated myeloid derived suppressor cells (MDSCs) in an allogeneic bone marrow transplant model (27). Whether ILC2_10_ specifically have protective functions in cell therapies for GVHD, and if ILC2_10_ would have any direct effects on allogeneic T cell responses, has not been explored. However, growing reports support ILC2_10_ could be harnessed to promote immune tolerance (9–14).

Several challenges have prevented assessment of the cell therapy potential of human ILC2_10_, the first and foremost being the relative paucity of human ILC2s in peripheral blood and lack of effective expansion protocols. To overcome this challenge, we developed methods to robustly expand ILC2_10_ that display a stable phenotype. We then demonstrate cell therapy with ILC2_10_ limits GVHD in a preclinical humanized mouse model, and that ILC2_10_ do not impair GVL responses critical for HSCT therapeutic effectiveness. We define novel markers of expanded ILC2_10_ that enable discrimination and isolation of ILC2_10_ from conventional ILC2s. We also identify ILC2_10_ directly suppress allogeneic T cell responses in a manner distinct from Tregs, as well as display distinct protective mechanism than what has been reported with conventional ILC2s. Findings herein provide proof-of-concept data to support cell therapy applications of human ILC2_10_ for T cell-driven pathologies such as those that underlie GVHD.

## Results

### Expansion of human ILC2s with an ILC2_10_ phenotype

Recent studies reporting protective functions of ILC2s and ILC2_10_ in multiple contexts supported that targeting or harnessing ILC2s in immunotherapy approaches may have benefits in dampening harmful inflammation. To explore this possibility, we first had to develop methods to isolate and expand large numbers of human ILC2s. ILC2s are present in very low abundance in peripheral blood and tissues, and most expansion protocols from mouse rely on *in vivo* cytokine administration prior to isolating ILC2s from bone marrow or other tissues, which would not be compatible with cell therapy approaches. Here, we isolated human ILC2s, as well as NK cells and ILC3s for comparison, from normal donor peripheral blood mononuclear cells (PBMCs) by flow cytometry sorting using a combination of ILC markers and gating out lineage positive cells (**Supplemental Figure.1A, B**). CD56^dim^ NK cells were sorted as live lineage^-^CD45^+^CD56^+^CD16^+^, ILC2s as live lineage^-^CD45^+^CD94^-^CD16^-^NKG2D^-^CD127^+^CRTh2^+^CCR6^-^ and ILC3s as live lineage^-^CD45^+^CD94^-^CD16^-^NKG2D^-^CD127^+^CRTh2^-^CD117^+^CCR6^+^. NK cells were expanded in NK MACS media supplemented with IL-2, IL-15 and IL-18, ILC2s were then expanded in X-VIVO media supplemented with IL-2, IL-7 and IL-33, and ILC3s were expanded in X-VIVO media with IL-1β, IL-2, IL-7 and IL-23. ILC2s and ILC3s exhibited robust expansion using this protocol, resulting in an average of 2.3×10^4^ ± 7.2×10^3^ and 9.8×10^3^ ± 3.6×10^3^ fold expansion respectively after 34 days in culture(**Supplemental Figure.1C**). Post-expansion, ILC2s strongly co-expressed IL-4, IL-13, IL-9 and GM-CSF following PMA-ionomycin stimulation, with low to no expression of IFN-γ, IL-17A or IL-22(**Figure.1A,B, Supplemental Figure.1D**). In contrast, expanded CD56^dim^ NK cells co-expressed IFN-γ and TNF-α and ILC3s expressed IL-22 and GM-CSF, with low IL-17A expression(**Figure.1A,B, Supplemental Figure.1D,2A**). Secreted cytokine production in expanded ILCs was quantified using cytometric bead array(CBA) following stimulation with IL-2. ILC2s secreted high amounts of IL-4, IL-9, IL-5 and IL-13, whereas NK cells and ILC3s did not produce significant amounts of these cytokines and instead produced canonical cytokines including IFN-γ and TNF-α or IL-22, respectively(**Figure.1C**, **Supplemental Figure.2B**). In addition to cytokine expression, we also examined chemokine production. Expanded ILC2s also produced the chemokines CCL2, CCL11, CCL20 and CXCL10 (**Supplemental Figure.2D**). To confirm ILC2 stability, we assessed the phenotype of expanded ILC2s when cultured in ILC3 or NK cell medium for 10 days(**Supplemental Figure.3A-J**). Expanded ILC2s maintained expression of signature cytokines and did not upregulate cytokines associated with NK cells or ILC3s, supporting once activated and expanded, human ILC2s from healthy donor blood maintained a stable ILC2 phenotype.

**Figure 1.**
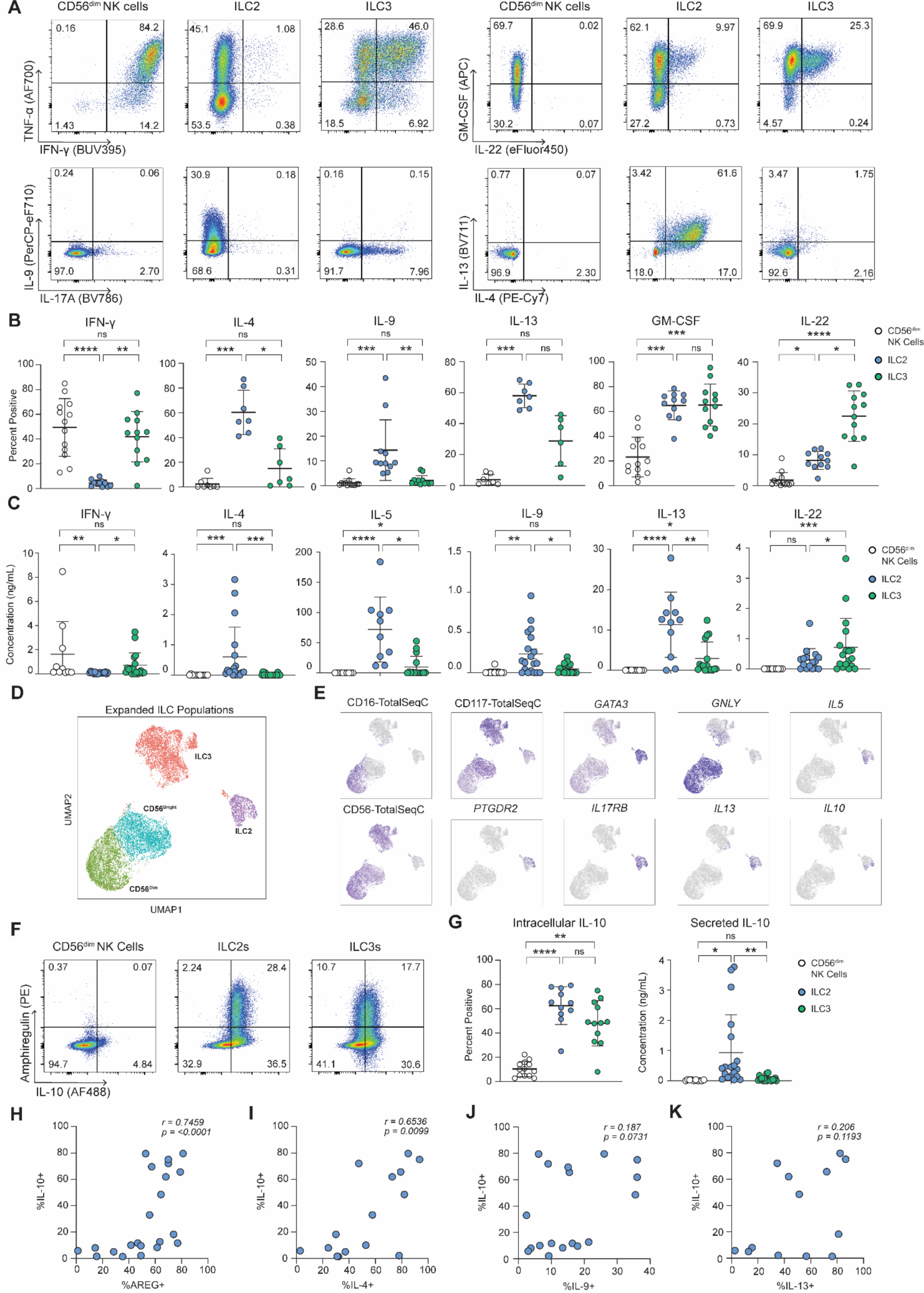
Expanded human ILC2_10_ maintain expression of signature cytokines. To explore the role of human ILC2s in GVHD, human ILCs were isolated from peripheral blood using flow cytometry and expanded *ex vivo* using ILC subset-specific cytokines, as ILC2s are present in very low abundance in peripheral blood. ILC2s expanded using our approach displayed and ILC2_10_ phenotype. Representative (**A**) and average (**B**) of intracellular cytokine staining for signature cytokines in CD56^dim^ NK cells, ILC2s and ILC3s at day 20 of expansion after stimulation with phorbol 12-myristate 13-acetate(PMA) and ionomycin. (**C)** Cytometric bead array analysis of secreted cytokines after plating ILCs at a concentration of 2×10^5^ cells/mL for 16-hour stimulation with 100U/mL of IL-2. (**D**) UMAP representation of CITE-seq of expanded ILC subsets. ILC subsets indicated were isolated by flow cytometry, expanded and then stained with CITE-seq antibody cocktail as outlined in **Supplemental Table 2**. Each population was also labeled with a unique hashtag antibody to facilitate post sequencing identification. (**E**) Feature plots showing protein level expression of CD16, CD56 and CD117 and RNA level expression of *PTGDR2, GATA3, IL17RB, GNLY, IL13, IL5* and *IL-10* on expanded ILC subsets. (**F**) Representative intracellular IL-10 and amphiregulin expression by expanded CD56^dim^ NK cells, ILC2s and ILC3s after PMA/Ionomycin stimulation. (**G**) Average intracellular IL-10 expression assessed by flow cytometry (n=13, 11 and 12 for CD56^dim^ NK cells, ILC2s and ILC3s respectively) and secreted IL-10 by cytometric bead array by expanded ILC subsets (n=12, 19 and 20 for CD56^dim^ NK cells, ILC2s and ILC3s respectively). Correlation of IL-10 expression with amphiregulin (**G**), IL-4 (**H**), IL-9 (**J**) or IL-13 (**K**)(n=22, 15, 18 and 13 respectively) on expanded ILC2s with ILC2_10_ phenotype.

To confirm expanded ILC2s maintained an ILC2-associated gene and protein expression profile, as well as enable detailed phenotyping of expanded ILC2s, we preformed Cellular Indexing of Transcriptomes and Epitopes by Sequencing(CITE-seq) on expanded ILCs. Expanded ILCs were stained with a TotalSeq-C antibody cocktail and labelled with hashtag antibodies to enable easy identification after sequencing(**Figure.1D, Supplemental Table 1**). All ILCs examined expressed lineage defining markers that differentiated them from other family members. CD56^bright^ and CD56^dim^ NK cells expressed *GNLY* and CD56, while the CD56^dim^ NK cells expressed CD16. ILC3s strongly expressed CD117 and ILC2s expressed *PTGDR2, GATA3, IL17RB, IL13* and *IL5* (**Figure.1E**). Intriguingly, one of the top genes identified as being uniquely expressed by *ex vivo* expanded ILC2s was *IL-10* (**Figure.1E**), which we validated by flow cytometry and cytometric bead array(**Figure.1F,G**). Expanded ILC2 supernatants had consistently high levels of IL-10 secretion across all donors, and a high proportion of IL-10^+^ cells were noted in the majority of donors examined(**Figure.1F,G**). Therefore, ILC2s isolated and expanded using our approach had a phenotype consistent with being ILC2_10_.

Due to recent studies that identified murine ILC2_10_ as having immunoregulatory properties, including being able to suppress T cells(10, 13, 33), we assessed expanded ILC2s with ILC2_10_ phenotype for potential immunoregulatory properties beyond IL-10 production, or expression markers that overlapped with other regulatory populations such as regulatory CD4^+^FOXP3^+^ Tregs and T regulatory 1 (Tr1) cells. Low or background expression of checkpoint molecules such as PD-1, PD-L1, CTLA-4, LAG3 was observed at both the RNA and protein level, and no *FOXP3* gene transcript was observed**(Supplemental Figure.4A,B)**(34, 35). Additionally, we did not detect *IL12A* and *EBI3*, which make up the regulatory cytokine IL-35**(Supplemental Figure.4A,B)**. Despite ubiquitous transcriptional expression of *TGFΒ1* across ILC subsets, no protein-level expression of active TGF-β1 was observed in expanded NK cells, ILC2s or ILC3s(**Supplemental Figure.2C, Supplemental Figure.4B**). While these regulatory molecules were absent, expanded ILC2s expressed *ENTPD1*, the gene for CD39 (**Supplemental Figure.4C**) and also weakly expressed *NT5E* (CD73). However only ILC2s and CD56^bright^ NK cells exhibited surface CD73 protein expression(**Supplemental Figure.4C**). Flow cytometry analysis verified expanded ILC2s co-expressing high levels of CD73, in line with reports of *ex vivo* ILC2s(36) as well as CD39(**Supplemental Figure.4D,E**), which are known to inhibit the functional activity of T cells(37, 38). IL-10 expression on ILC2s was correlated with co-expression of IL-4, as well as amphiregulin (AREG) but had no significant correlation to expression levels IL-9 or IL-13 cytokines(**Figure1.H-K**). High amphiregulin was also notable, as it is a key tissue reparative effects(27, 39, 40), and was linked to ILC2s protective effects in limiting GVHD, supporting dual regulatory and reparative potential of expaned ILC2_10_.

### Adoptive transfer of allogeneic human ILC2_10_ suppresses xenogeneic GVHD

Pre-clinical models and patient-based studies have linked CD4^+^ T helper 1(Th1) and CD8^+^ cytotoxic T cells (Tc1) to aGVHD pathology(41–44). 30% of aGVHD patients do not respond to first line corticosteroid treatment, with less than one-third of non-responders surviving past one year(45), highlighting the need to identify cells and molecules that limit pathogenic T cell responses post-HSCT. Due to prior mouse studies supporting potential applications of ILC2 transfer for GVHD(27, 46), and increasing appreciation of immunoregulatory functions of ILC2_10_, we sought to assess the cell therapy potential of human ILC2_10_ using a preclinical xenogeneic GVHD model(xenoGVHD). In this model, 6– 8-week-old female NOD-scid-IL2Rg^null^ (NSG) mice are irradiated and injected with human PBMCs. T cells from transferred PBMCs engraft into NSG mice and induce tissue pathology similar to acute GVHD (47). NSG mice are monitored for signs of xenoGVHD, using weight-loss and a composite xenoGVHD score, which measures fur loss, skin inflammation, hunch, activity, and pain (**Figure.2A**). T cell engraftment, as well as detailed phenotyping is performed on circulating immune cells and immune cells within tissues at endpoint or indicated times.

Using this model, we assessed whether cell therapy with ILC2_10_ could limit human T cell-mediated pathology. NSG mice that received PBMCs alone or PBMCs and ILC2_10_ were first compared for signs of GVHD and how this corresponded with T cell engraftment. After 14 days, engraftment of human CD45^+^CD3^+^T cells was observed in the blood, with no significant difference in T cell proportions between NSG mice given human PBMCs alone or mice treated with ILC2_10_, indicating ILC2_10_ did not inhibit T cell engraftment(**Figure.2B-D**). This was of note, as cell therapy with Tregs in this xenoGVHD model results in impaired T cell engraftment (48). Despite not impacting T cell engraftment, female NSG mice treated with ILC2_10_ had significantly delayed onset and reduced GVHD symptoms, including decreased weight-loss across 3 independent experiments (**Figure.2E-H**). These effects were also observed in separate experiments in male NSG mice, although with slight differences in xenoGVHD kinetics(**Supplemental Figure.5A-C**). This single infusion of ILC2_10_ not only reduced xenoGVHD severity but also significantly improved overall survival(**Figure.2I**). Notably ILC2_10_-treated mice only reached endpoint due to weight-loss, but other GVHD symptoms were still very low and mice would otherwise not have reached endpoint(**Figure.2E,F**).

To assess if ILC2_10_ could also treat GVHD upon onset, allogeneic ILC2_10_ were adoptively transferred after T cell engraftment and onset of xenoGVHD symptoms. Here, T cell engraftment was confirmed prior to ILC2_10_ transfer, and mice had begun displaying early xenoGVHD symptoms. When human ILC2_10_ were transferred at day 9, this resulted in similar protection from xenoGVHD to when ILC2_10_ were administered prophylactically at day 0(**Supplemental Figure 6**). Taken together, cell therapy with allogeneic expanded human ILC2_10_ limited pathology and enhanced overall survival in this model of GVHD when administered both prophylactically and upon GVHD onset.

### ILC2_10_ inhibit allogeneic T cell responses in xenoGVHD

We next sought to determine how ILC2_10_ protected from GVHD. Similar to previous reports, T cells made up >96% of engrafted human CD45^+^cells across all organs analyzed (**Figure 2B-D** and **Supplemental Figure.7A-D**)(47). Despite no differences in engraftment at day 14, we noted at endpoint a moderate increase in CD4^+^ Th cells and reciprocal decrease in cytotoxic CD8^+^ T cells in the blood, bone marrow and spleen of NSG mice receiving ILC2_10_ cell therapy(**Figure.2J**). We observed CD4^+^ and CD8^+^ T cells had reduced expression of Ki-67 in the blood, bone marrow and spleen of NSG mice treated with ILC2_10_, indicating reduced T cell proliferation (**Figure.3A-D**, **Supplemental Figure.8A,B**). This was not accompanied by changes in T cell expression of checkpoint molecules examined including PD-1, CTLA-4 and CD25 (**Supplemental Figure.9A-C**).

**Figure 2.**
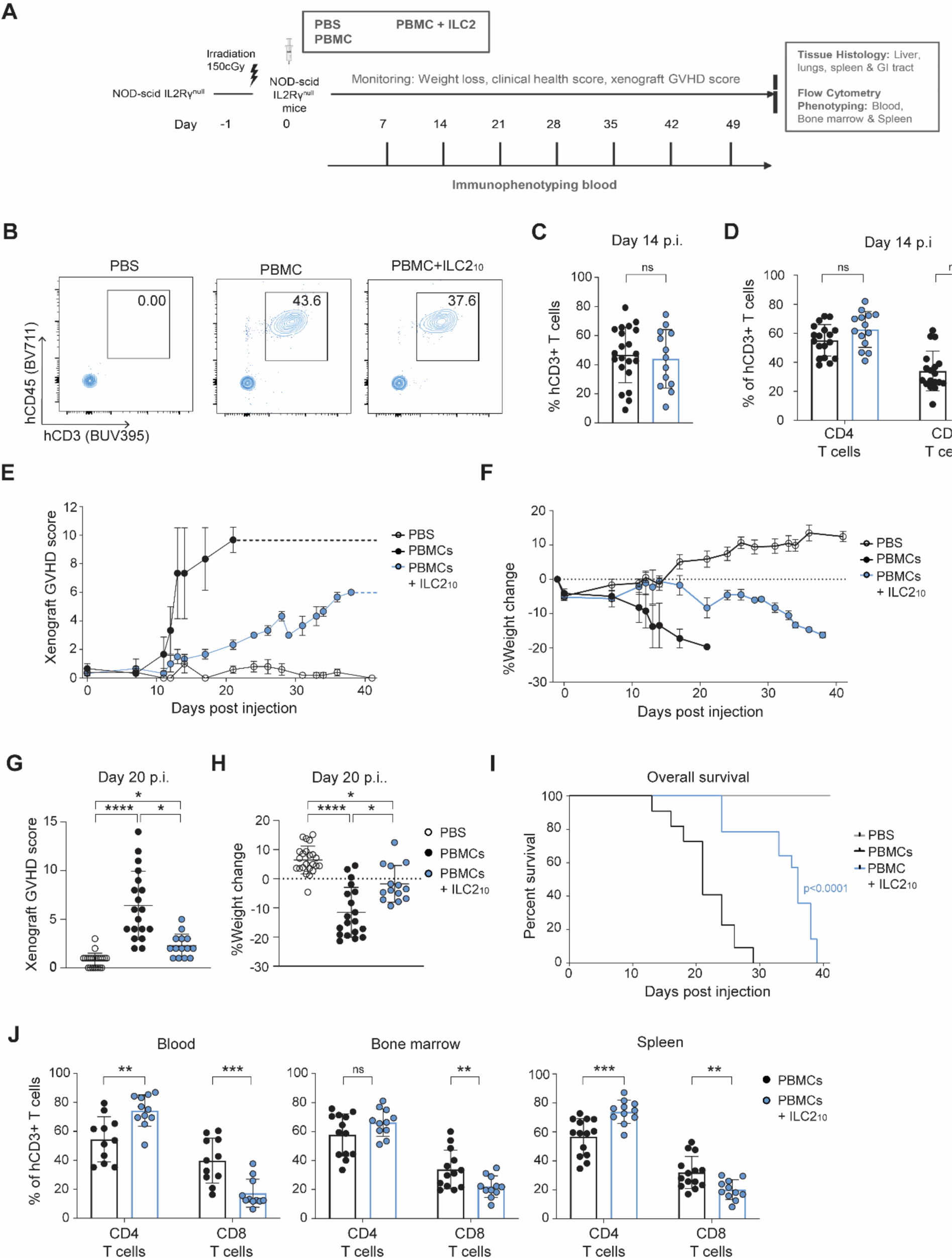
Cell therapy with human ILC2_10_ suppresses xenogeneic GVHD. To assess whether human ILC2_10_ could limit GVHD, we employed an established xenogeneic model of GVHD involving PBMC transfer into NOD-scid-IL2Rg^null^ mice (NSG). (**A**) Overview of the xenogeneic GVHD model to assess whether ILC2_10_ would prophylactically limit GVHD. Briefly, NSG mice are irradiated and receive either PBS, human PBMCs, to induce multiorgan tissue pathology, or PBMCs and ILC2_10_ together. Mice are monitored for symptoms of xenogeneic GVHD including weight loss, survival and a composite xenogeneic GVHD score consisting of hunch, skin inflammation and fur loss, activity, pain and percent weight change. Blood is drawn at 7-day intervals for confirmation of human immune cell engraftment and immunophenotyping. At the humane or experimental endpoint, mice are euthanized, and tissues collected for histology or flow cytometry analysis. (**B**)Representative engraftment human CD3^+^ CD45^+^ T cells at day 14 post injection of human PBMCs alone or with ILC2_10_. **C**,**D** Average engraftment of human CD3^+^ CD45^+^ (**C**), or CD4^+^ and CD8^+^ T cells (**D**), in the blood 14 days post injection across 3 independent experiments (n=21 for NSG mice receiving human PBMC and n=13 for NSG mice receiving human PBMC plus allogeneic ILC2_10_ mice from 3 independent experiments). **E,F** (**E**) Representative xenogeneic GVHD score (n=3/group in this experiment), and (**F**) representative weight loss (n=3/group in this experiment) of 3 independent experiments. **G,H** Average of xenogeneic (**G**) GVHD score and (**H**) weight loss of mice at day 20 post injection across three independent experiments (n=23 for PBS treated NSG mice, n=19 for NSG mice receiving human PBMCs alone and n=14 for NSG mice receiving human PBMC plus allogeneic ILC2_10_ respectively across 3 independent experiments). (**I**) Survival of mice treated with ILC2_10_ across 3 independent experiments (n=23, 19 and 14 for NSG mice receiving PBS, PBMC and PBMC+ ILC2_10_ respectively). Experimental endpoint was day 40 for any mice not reaching human endpoint. (**J**) Average proportion of CD4^+^ and CD8^+^ T cells of total human CD3^+^ T cells in the blood, bone marrow and spleens of NSG mice at endpoint (n=11 and 13 from PBMC and PBMC+ ILC2_10_ respectively from 3 independent experiments).

**Figure 3.**
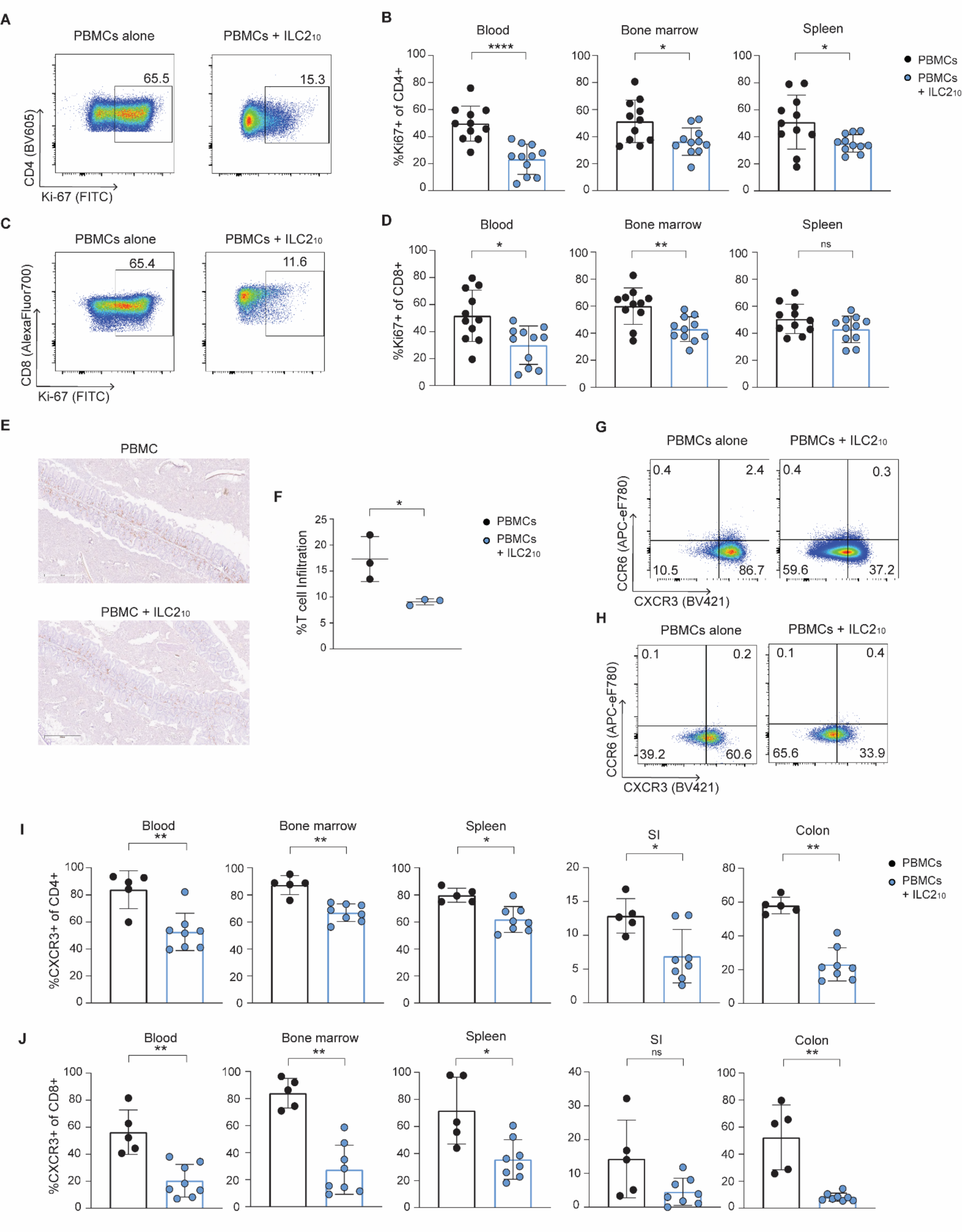
Cell therapy with ILC2_10_ suppresses CD4^+^ and CD8^+^ T cell proliferation, intestinal infiltration and CXCR3 expression. Effect of ILC2_10_ transfer on T cell phenotypes and proliferation in xenogeneic GVHD model were assessed by flow cytometry at endpoint. (**A**) Representative Ki-67 expression by circulating human CD4^+^ T cells from the blood at end point (**B**) Average Ki-67 expression in blood, bone marrow or spleen at endpoint by human CD4^+^ T cells from 3 independent experiments(n=11 mice per condition from 3 independent experiments). (**C**) Representative Ki-67 expression by circulating human CD8^+^ T cells. (**D**) Average Ki-67 expression in the blood, bone marrow or spleen at endpoint on human CD8^+^ T cells from 3 independent experiments(n=11 mice per condition from 3 independent experiments) (**E**) Representative human CD3 immunohistochemistry of the colon of mice treated with PBMCs or PBMCs plus ILC2_10_ (10x magnification). (**F**) Average T cell infiltration in the colon of mice treated with PBS, PBMCs or PBMCs + ILC2_10_. Quantified using CD3 immunohistochemistry and the HALO algorithm and normalized to background in PBS mice. **G,H** Representative CXCR3 expression on circulating human CD4^+^ (**G**) or CD8^+^ (**H**) T cells. Average CXCR3 expression on human CD4^+^ (**I**) or CD8^+^ (**j**) T cells in the blood, bone marrow, spleen, colon, and small intestines at experimental endpoint (n=5 from 2 independent experiments).

Previous research has indicated that CXCR3^+^T cells drive trafficking to target organs including intestines in mouse models of GVHD(49, 50). CXCR3 also marks CD4^+^ Th1 and CD8^+^ Tc1 cells(51–53) which induce intestinal damage in GVHD models and correlate with disease in patients(41, 43). We therefore asked whether ILC2_10_ influenced T cell phenotypes and intestinal trafficking within the xenoGVHD model. CD3 immunohistochemical(IHC) staining of tissues revealed a stark reduction in T cell infiltration into colons of ILC2_10_-treated mice, with no change in T cell infiltration noted in the lungs, spleen, or liver(**Figure.3E,F**, **Supplemental Figure.10A-G**). Across compartments examined, mice receiving ILC2_10_ cell therapy had reduced CXCR3 expression by CD4^+^ and CD8^+^ T cells(**Figure.3G-J**, **Supplemental Figure.8C,D**), with no increase in CRTh2 or CCR6, markers of CD4^+^ T helper 2 (Th2) and T helper 17 (Th17) cells, respectively(52–54). Adoptive transfer of human ILC2_10_ therefore did not promote differentiation to other T cell subsets, but inhibited CD4^+^Th1 and CD8^+^Tc1 cells(**Figure.3G-J, Supplemental Figure.8C,D**)(52–54).

### Inverse correlation between ILC2s and pathogenic T cells in HSCT recipients

Due to ILC2_10_-mediated suppression of CD4^+^ Th1 and CD8^+^ Tc1 cells in the xenoGVHD model, and prior studies reported increased CD56^bright^ NK cells, ILC2s and ILC3s in peripheral blood of HSCT patients that did not develop GVHD compared to those that did(32), we were interested in the relationships between ILCs and T cell subsets in HSCT recipients with differing clinical outcomes. We analyzed the proportion of subsets of ILCs and T cells in the blood of HSCT patients at the time of aGVHD diagnosis and compared proportions of these populations in patients without aGVHD and healthy donors(**Supplemental Figure.11A**, **Supplemental Table 2**). All HSCT recipients were given prophylactic post-transplant cyclophosphamide, and two were treated with additional immunosuppression at the time of sample acquisition. In line with previous literature, patients without aGVHD had decreased proportions of CD56^dim^ NK cells and increased proportions of CD56^bright^ NK cells(30, 31)(**Supplemental Figure.11B,C**). Similarly, patients that developed aGVHD had a reduced proportion of total helper ILCs (Lin^-^ CD127^+^)(**Figure.4A,B**), with reduced proportions of ILC1s, ILC2s and ILC3s compared to healthy controls(**Figure.4C,D**). HSCT patients without aGVHD, however, had significantly higher proportions of ILC2s, but no significant differences in ILC1s or ILC3s compared to patients with aGVHD with our sample size (**Figure.4C,D**).

**Figure 4.**
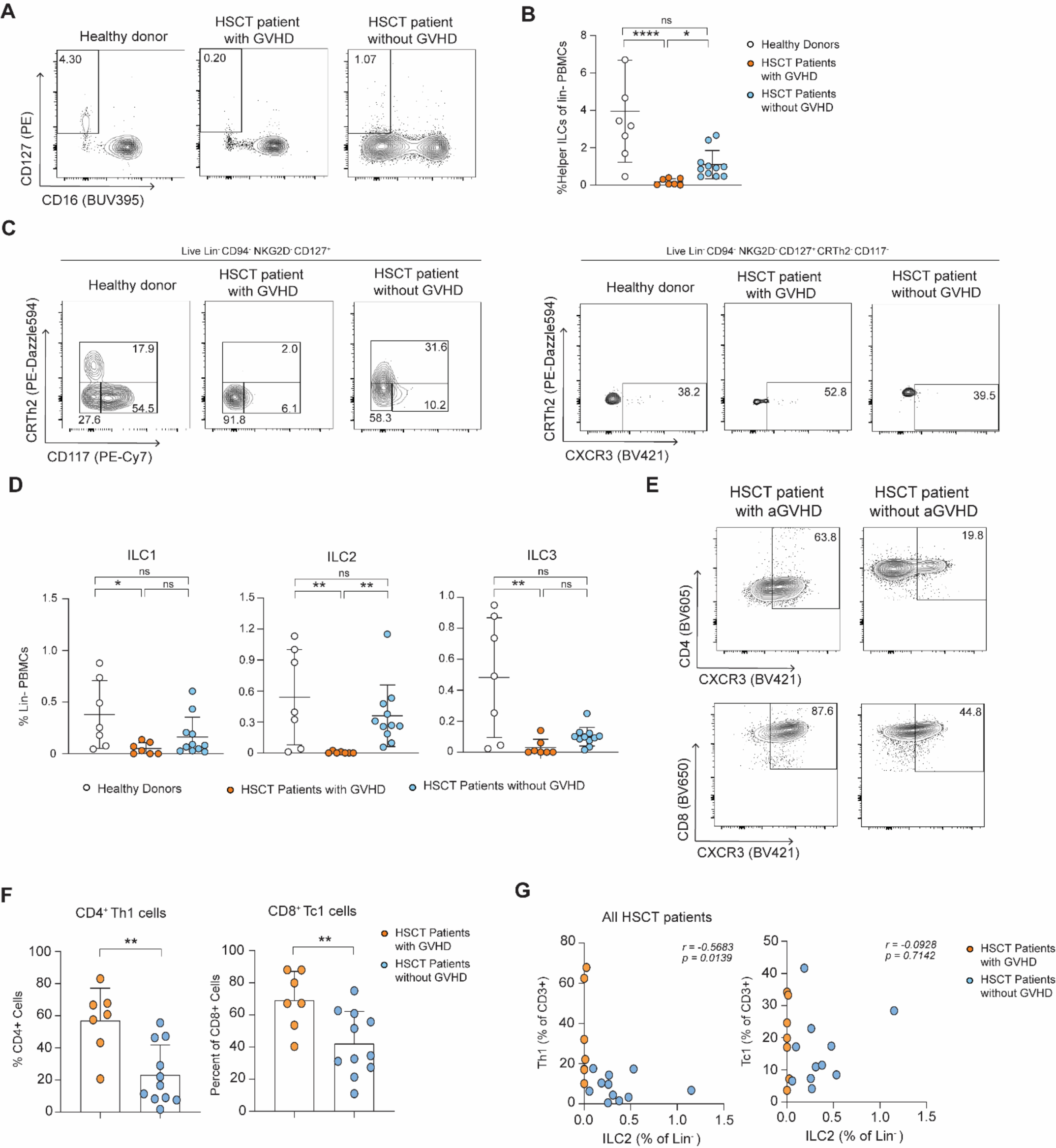
Increased circulating ILC2s is associated with decreased CD4^+^ Th1 cells in HSCT recipients protected from GVHD. PBMCs from healthy donors, and HSCT patients with or without GVHD were assessed for ILC proportions and phenotypes using flow cytometry. **A,B** Representative flow plots (**A**) and summary graphs (**B**) of total CD127^+^ helper ILCs in PBMCs as a proportion of live CD45^+^ lineage negative cells. Lineage markers are CD3, CD4, CD8, CD14, CD19, CD20, CD33, CD34, CD123, CD138, CD303, FCεRI and TCR𝛾𝛿. **C,D** Representative (**C**) and average (**D**) CD117^-^, CRTh2^-^, CXCR3^+^ ILC1s, CRTh2^+^ ILC2s and CRTh2^-^ CD117^+^ ILC3s in PBMCs as a proportion of live CD45^+^ lineage negative cells. (**E**) Representative CD4^+^ Th1 and CD8^+^ Tc1 cells in HSCT patients with or without aGVHD development. (**F**) CD4^+^ Th1 and CD8^+^ Tc1 cells across HSCT patients as a proportion of CD4^+^ and CD8^+^ T cells respectively. (**G**) Correlation of Th1 and Tc1 cells with ILC2s across all HSCT patients.

We next examined the correlation of ILCs with T cell subsets in HSCT recipient peripheral blood. Similar to what we observed with ILC2 therapy in the xenoGVHD model, HSCT recipients with aGVHD had increased CD4^+^ Th1 and CD8^+^ Tc1 cells as a proportion of total CD4^+^ and CD8^+^ T cells respectively(**Figure.4E,F**), with no significant difference in circulating Th17 cells (51–53). Elevated ILC2s, significantly correlated with low proportions of Th1 cells as a percent of CD3^+^ T across all HSCT patients(**Figure.4G-H**). Therefore, ILC2s proportions following HSCT are associated with decreased pathogenic T cell responses, mirroring findings in our xenoGVHD model, and further supporting the therapeutic potential of ILC2s for GVHD. While this paralleled the inverse relationship observed in xenoGVHD model, a caveat was that we were not able to specifically examine whether these ILC2s were specifically ILC2_10_, as markers of IL10^+^ILC2s had not been established.

### Expanded IL-10^+^ILC2s regulate CD4^+^ and CD8^+^ T cells via IL-4 and IL-10

We next asked whether ILC2_10_ could directly regulate allogeneic T cells. *In vitro* co-cultures were performed with ILC2_10_ and allogeneic naïve CD4^+^ or CD8^+^ T cells. Culture with ILC2_10_ resulted in a significant reduction in IFN-γ^+^ and IFN-γ^+^/TNF-α^+^ CD4^+^ T cells(**Figure.5A,B**). Decreased IFN-γ was also observed when allogeneic ILC2_10_ were cultured with CD8^+^ T cells(**Figure.5C,D**). To confirm *in vivo* findings that cell therapy with ILC2_10_ reduced Th1 and Tc1 cells without increasing proportions of Th2 or Th17 cells, we also examined cytokines expressed by other T cell subsets. No increase in IL-17A, IL-22, IL-4, IL-9 or IL-13 was observed with addition of ILC2_10_ to cultures with CD4^+^ or CD8^+^ T cells(**Supplemental Figure.12A,B**). Thus, decreased IFN-γ does not appear to be due to T cell differentiation into other T cell subsets, but ILC2_10_-mediated inhibition of CD4^+^ Th1 and CD8^+^Tc1 cells.

**Figure 5.**
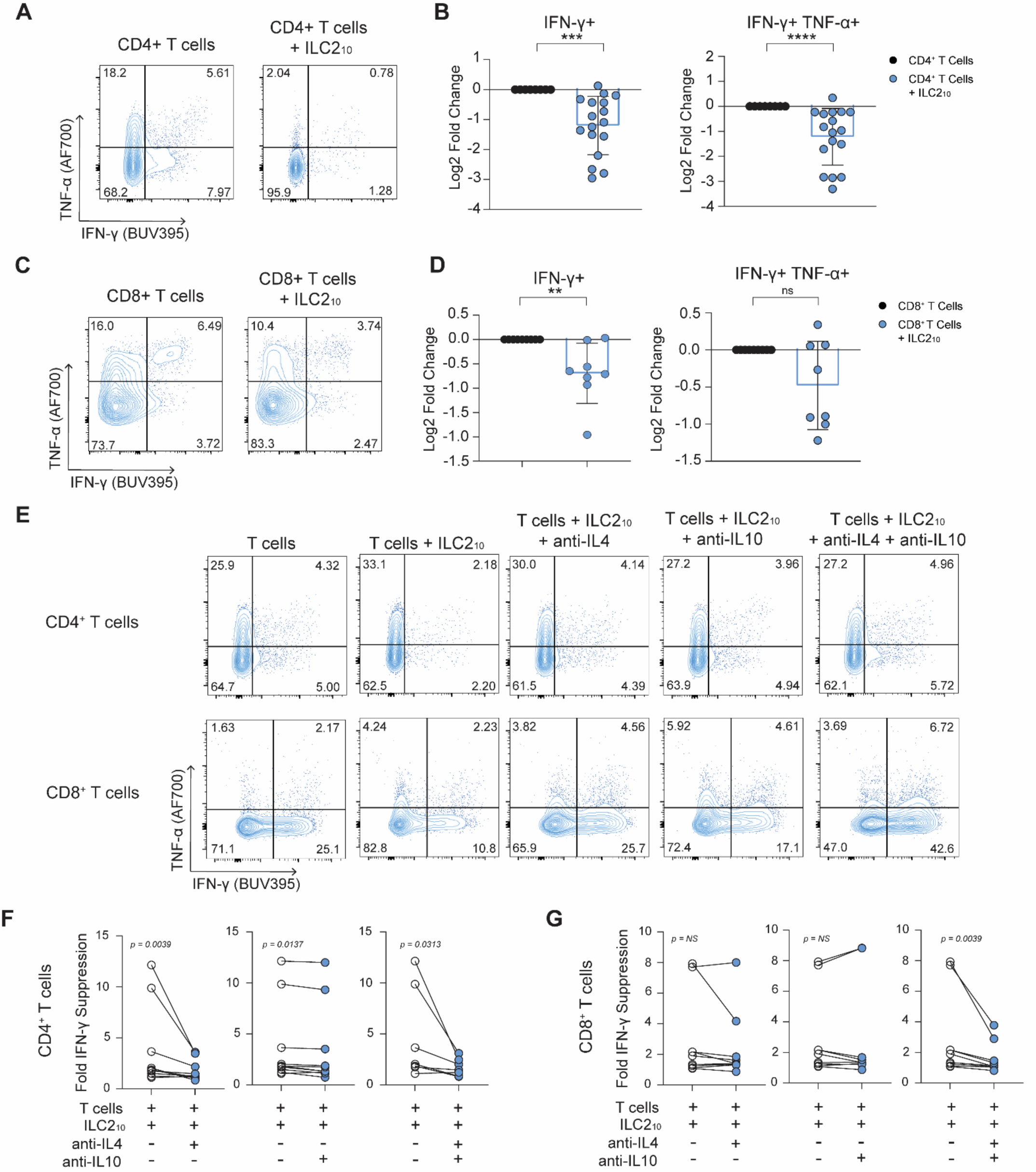
ILC2_10_ suppress CD4^+^ and CD8^+^ T cell cytokine production via a combination of IL-4 and IL-10. To determine whether cell therapy with ILC2_10_ protective effects were due to direct effects of ILC2_10_on allogeneic CD4^+^ and CD8^+^ T cells, we performed *in vitro* co-cultures. Expanded ILC2_10_ were cultured with naïve CD4^+^ and CD8^+^ T cells that were activated with anti-CD3/anti-CD28 beads. After 4 days in culture, intracellular cytokine staining was assessed by flow cytometry. (**A**) Representative IFN-γ and TNF-α expression by naïve CD4^+^ T cells cultured with or without allogeneic ILC2_10_ at day 4. (**B**) Average decrease in IFN-ψ production or IFN-ψ and TNF-α co-expression represented as log_2_fold change compared to CD4^+^ T cells alone (n=17). (**C**) Representative IFN-γ and TNF-α expression on CD8^+^ T cells co-cultured with allogeneic ILC2_10_ and stimulated with anti-CD3/CD28 beads over 4 days by representative flow cytometry plots. (**D**) Average decrease in IFN-ψ and co-expression of IFN-ψ and TNF-α representative as log_2_fold change compared to CD8^+^ T cells alone (n=8). (**E**) Representative CD4^+^ and CD8^+^ T cell IFN-ψ expression when cultured with or without ILC2_10_ in the presence of indicated blocking anti-IL-4 and/or anti-IL-10 antibodies. Effects on fold change in IFN-γ expression by CD4^+^ (**F**) or CD8^+^ (**G**) T cells after culture with ILC2_10_ with the addition of anti-IL-4 (n=10), anti-IL-10 (n = 10) or anti-IL-4 and anti-IL-10 (n=7) blocking antibodies

We next explored whether various molecules associated with ILC2s or ILC2_10_ identified by CITE-seq controlled ILC2-mediated regulation of T cells interactions. We noted that allogeneic T cells activated in the presence of supernatants from IL-2-stimulated ILC2_10_ had comparable suppression of T cells to that of adding ILC2s to T cell cultures directly(**Supplemental Figure.12C,D**), indicating one or more secreted factors produced by ILC2_10_ underlies their ability to inhibit T cells.

ILC2_10_ were therefore cultured with allogeneic T cells in the presence or absence of neutralizing antibodies or inhibitors to secreted molecules of interest(IL-10, CD39, CD73), as well as ILC2 cytokines (IL-4, IL-9, IL-13)(6, 55, 56). The addition of an IL-4-neutralizing antibody strongly reduced ILC2_10_-mediated suppression of IFN-γ by CD4^+^ T cells *in vitro,* but blocking IL-9, IL-13 or CD39/CD73 had no effect(**Figure.5E,F**, **Supplemental Figure.12E, Supplemental Figure.13A**). Neutralizing IL-10 also decreased suppression of CD4^+^ T cell-IFN-γ, but to a lesser extent than IL-4(**Figure.5E,F, Supplemental Figure.13A**). In contrast, the combination of anti-IL-4 and anti-IL-10 abrogated ILC2_10_-mediated suppression of IFN-γ by CD8^+^ T cells, which was not observed with anti-IL-4 or anti-IL-10 alone(**Figure.5E,G, Supplemental Figure.13B**). ILC2_10_ suppression of allogeneic CD4^+^ and CD8^+^ T cells cytokine production requires IL-4 and IL-10, with differing contributions of IL-4 and IL-10 to the regulation of CD8^+^ and CD4^+^ T cells.

### Allogeneic human ILC2_10_ do not suppress T cell mediated graft-versus-leukemia effects

The GVL effect is critical for success of allogeneic HSCT for treating hematological malignancies. While cell therapy with ILC2_10_ did not impair T cell engraftment, ILC2_10_ suppressed xenogeneic T cell responses. A key question therefore was whether cell therapy with ILC2_10_ would inhibit T cell-mediated GVL effects. Within our clinical cohort, we assessed circulating ILC2 proportions and how they correlated with relapse status. No significant differences in ILC2 proportions was observed in the blood of patients which went on to experience relapse of their malignancy, supporting that increased proportions of ILC2s are not associated with reduced GVL effects(**Figure.6A**).

We next assessed whether ILC2_10_ cell therapy would negatively impact T cell-mediated GVL-effects in a in a humanized mouse model. Briefly, MV4-11 cells, an acute myeloid leukemia cell line, were injected into mice 5 days prior to transfer of PBMCs alone or PBMCs with allogeneic expanded human ILC2_10_ (**Figure.6B**). The ability of T cells to kill MV4-11 cells was then assessed at day 14 following PBMC transfer. As expected, PBMC injection resulted in a reduction in MV4-11 cells in the bone marrow compared to mice receiving MV4-11 cells alone(**Figure.6C,D**). Mice treated allogeneic ILC2_10_ had comparable reductions in MV4-11 cells within the bone marrow to that observed with mice receiving PBMCs alone(**Figure.6C,D**), supporting ILC2_10_ were not impeding T cell-mediated GVL responses. Importantly, within the same mice, ILC2_10_ treatment reduced xenoGVHD symptoms similar to what was observed without MV4-11 transfer(**Figure.6E**), clearly showing simultaneous protection from pathogenic T cell responses that underlie GVHD without impairing T cell-mediated GVL effects. Collectively our findings support ILC2_10_ do not limit the anti-leukemic effect of allogeneic T cells despite protecting from GVHD, therefore representing a strong candidate for cell-based therapies for GVHD following HSCT.

**Figure 6.**
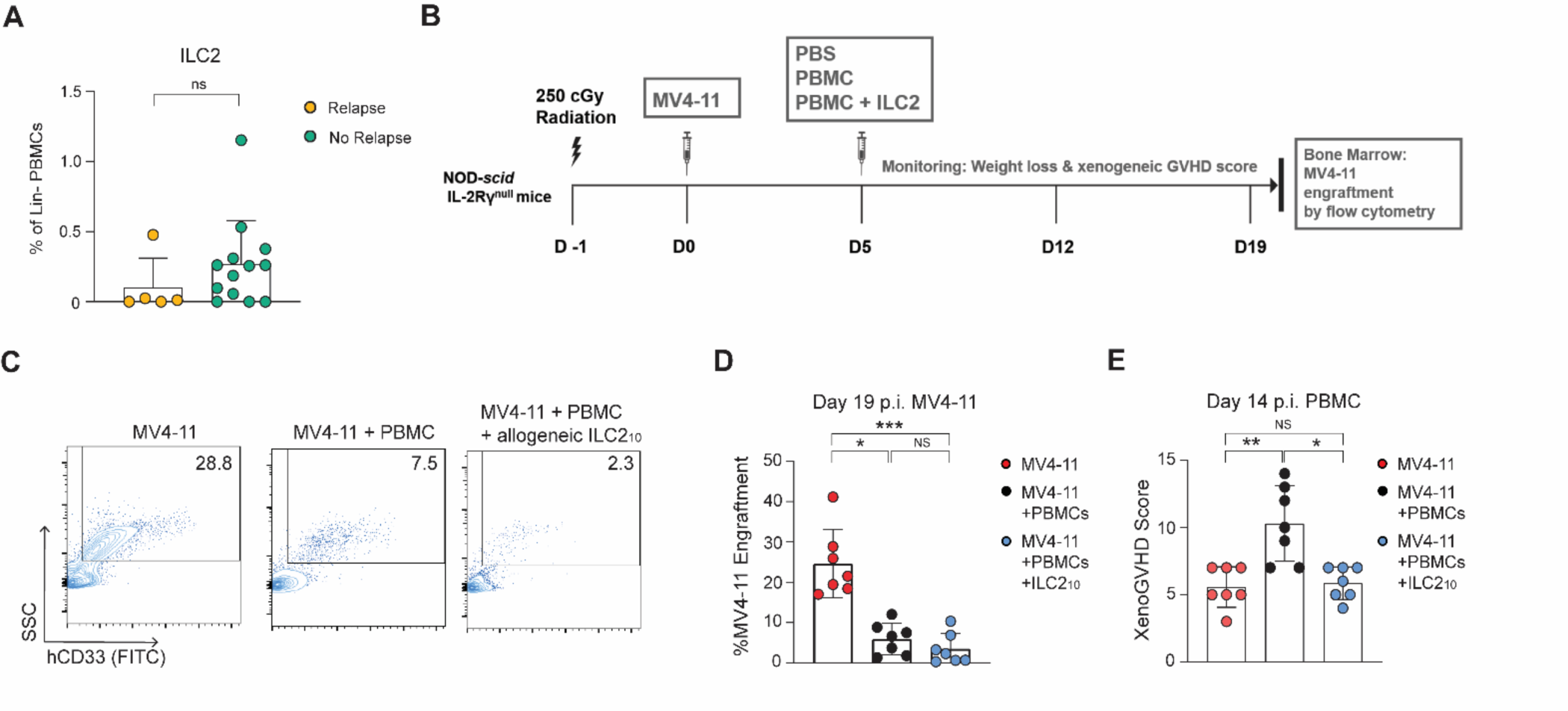
Cell therapy with ILC2_10_ does not impede T cell-mediated graft-versus-leukemia effect. **(A)** Circulating ILC2s from HSCT patients as a proportion of lineage negative PBMCs. Patients were grouped based on whether they experienced cancer relapse (n=5) or not (n=13). Follow up periods at time of assessment ranged from 30 to 120 days. (**B**) Overview of humanized GVL model, whereby MV4-11 AML cells are transplanted into NSG mice. PBMCs are administered at D5 following MV4-11 transfer, and engrafted T cells reduced MV4-11 cell engraftment. (**C**) Representative MV4-11 cells in bone marrow of NSG mice treated with PBS, PBMCs or PBMCs with ILC2_10_. (**D**) Average MV4-11 engraftment in NSG mice treated with PBS, PBMCs or PBMCs with ILC2_10_ from two independent experiments (n=7/group) (**E**) Average xenogeneic GVHD score at day 19 from 2 independent experiments (n = 7/group). Day 19 corresponded to 14 days following injection of PBMCs with or without ILC2_10_.

### *Ex vivo* expanded ILC2_10_ highly express CD49d and CD86

Despite consistently high levels of IL-10 in our *ex vivo* expanded ILC2_10_, we noted heterogeneity in the proportions of IL-10^+^ cells between ILC2 isolations from different donors. In order to improve consistency of ILC2_10_ in cell expansions, as well as define key molecules that differentiate IL-10^+^ILC2s from IL10^-^ILC2s, we performed differential gene expression analysis of *IL10*^+^ILC2s to *IL10*^-^ILC2s in our CITEseq dataset on expanded human ILCs. Elevated expression of CD86 and *ITGA4* (CD49d) was noted on the *IL10*^+^ILC2s(**Figure.7A**), which was confirmed by flow cytometry to identify IL-10^+^ILC2s (**Figure.7B,C**). We also examined CD117 expression, as *ex vivo* ILC2s had variable expression of CD117 during isolation, and KLRG1, which was previously reported to be expressed by IL-10-producing ILC2(13). Both IL-10^+^ and IL-10^-^ILC2s had very low expression of these molecules, although we noted a small but significant fold increase in CD117 expression(**Figure.7D,E**). Using tSNE clustering following flow cytometry staining, a distinct cluster expressing IL-10 was defined by high expression of CD49d and CD86 (**Figure.7F**). Dividing expanded ILC2s into populations based on expression of CD49d, CD86 or co-expression revealed CD49d and CD86 marked IL10-expressing ILC2s with the highest IL-10 expression on CD49d^+^CD86^+^ ILC2s(**Figure.7F**). CD49d and CD86 expression therefore identify *ex vivo* expanded ILC2_10_, and can be used to isolate ILC2_10_ in expansion cultures.

**Figure 7.**
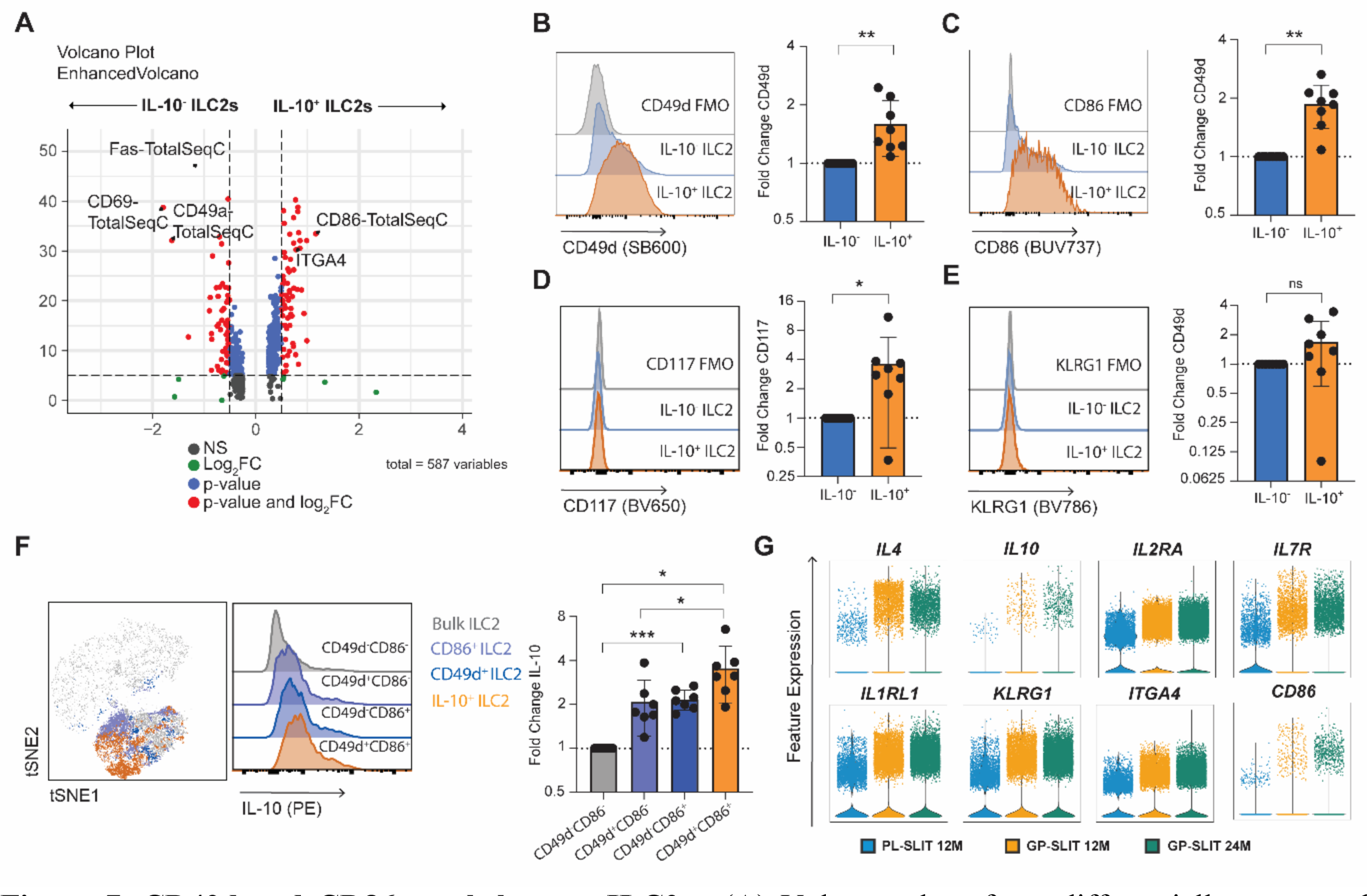
CD49d and CD86 mark human ILC2_10_. (**A**) Volcano plot of top differentially expressed transcripts and antibody-derived tags (ADT) between IL-10^+^ and IL-10^-^ expanded human ILC2s. To validate uniquely expressed surface markers of ILC2_10_, flow cytometry was performed on additional donors for CD49d CD86. **B-E**, Representative and average expression of CD49d (**B**), CD86 (**C**), as well as CD117 (**D**), and KLRG1 (**E**) by IL-10^+^ and IL-10^-^ ILC2s following PMA/ionomycin stimulation (n=7). (**F**) Average expression of IL-10 following PMA-ionomycin stimulation on ILC2s based on expression of CD49d and CD86 as a representative IL-10 expression, tSNE clustering based on co-expression and average graphs (n=7). (**G**) Violin plots of gene expression of CD86 and CD49d, as well as other ILC2_10_-associated molecules, from publicly available scRNAseq dataset of grass-pollen allergy patients treated with placebo (PL-SLIT) or grass-pollen allergen immunotherapy (GP-SLIT) at 12 or 24 month follow-up (n = 2 patients per group).

To assess the potential utility of these markers to identify ILC2_10_ in patient-based studies, we examined the publicly available scRNA-seq dataset of ILCs from patients with grass-pollen allergy treated with either grass-pollen sublingual allergen-specific immunotherapy(GP-SLIT) or placebo (PL-SLIT)(13). Here, the authors reported that patients treated with GP-SLIT had improved outcomes and increased proportions of ILC2_10_ at 12-month and 24-month follow-ups compared to placebo treated patients.

Further, the authors reported KLRG1 expression defined ILC2_10_, and pathways related to IL-10 signalling and regulation were upregulated in at both time points following GP-SLIT. We re-analyzed this scRNA-seq dataset to evaluate whether the increased IL-10 expression by ILC2s was associated with increases in CD49d and CD86. Following clustering of ILC2s from GP-SLIT treated patients, there was increased expression of *IL10* as reported, as well as *IL4* (**Figure.7G**). Further, GP-SLIT treated patients had elevated expression of *IL2RA, IL7R* and *IL1RL1,* receptors for the cytokines used for our ILC2_10_ expansion (**Figure.7G**). Notably, ILC2s from GP-SLIT treated patients had elevated expression of *ITGA4* (CD49d) and *CD86,* in addition to elevated *KLRG1* expression as reported (**Figure.7G**). This elevated expression of CD49d and CD86 that correlated with ILC2_10_ supports the potential broad applications of these markers to differentiate ILC2_10_ from inflammatory ILC2s in human studies.

## Discussion

This proof-of-concept study demonstrates the cell therapy potential of human ILC2_10,_ and their capacity to directly regulate harmful T cell responses that drive GVHD pathology. ILC2_10_ were robustly expanded using our approach, and displayed a stable phenotype. Importantly, ILC2_10_ cell therapy inhibited development of GVHD in humanized mouse models without impairing GVL effects essential for the success of HSCT. These findings mirror HSCT patient-based studies, where high ILC2 proportions are associated with protection from GVHD, without increased risk of cancer relapse. Human CD4^+^ and CD8^+^ T cells had reduced proliferation, tissue trafficking, and intestinal infiltration following ILC2_10_ treatment, and there were decreased proportions of CD4^+^Th1 and CD8^+^Tc1 cells across multiple tissue sites with ILC2_10_ therapy. Direct regulation of allogeneic T cell responses in this manner is unique to ILC2_10_ cell therapy, as it was not reported in mouse studies of conventional ILC2s, or in Treg studies using this xenogeneic GVHD model (27). CD86 and CD49d marked *ex vivo* expanded ILC2_10_, which suppressed IFN-γ in CD4^+^ and CD8^+^ T cells through a combination of IL-10 and IL-4. Altogether, our findings support that human ILC2_10_ have potential applications in cell therapies aimed at limiting immune-driven pathologies.

ILC2s display significant heterogeneity and can adopt proinflammatory or tissue protective functions depending on cytokines and microenvironment factors (2, 9, 10, 12–14, 33). ILC2_10_ have been shown to inhibit immune responses in diverse contexts, including inflammation in the lungs, intestines, and neurological system(9–14). In humans, individuals with grass-pollen allergies have ILC2s with reduced capacity to produce IL-10 following stimulation, and patients who respond to allergy immunotherapy have increased proportions of IL-10^+^ILC2s(13). In the context of transplantation, Huang *et al.* demonstrated mouse ILC2_10_ could prolong islet allograft survival by suppressing T cell attack of transplanted islets(33). These growing reports demonstrate important functions for ILC2_10_ in regulating immunity. However, to date we have lacked markers that accurately identify ILC2_10_ from conventional ILC2s. In one of the few studies that attempted to identify unique markers of ILC2_10_, Golebski *et al.* reported KLRG1^+^ILC2s produced IL-10 along with other signature ILC2 cytokines(13). However, KLRG1 has also been associated with inflammatory ILC2s, and may instead mark activated or memory like ILC2 populations, not specifically ILC2_10_(15–18). In our study, CITE-seq analysis of expanded human ILC2_10_ revealed limited expression of KLRG1, but instead high levels of CD49d and CD86. Flow cytometry confirmed that ILC2s co-expressing CD49d and CD86 had the greatest expression of IL-10.

To assess the potential broad utility of these markers, we reanalyzed the publicly available dataset from Golebski *et al* of ILCs isolated from grass-pollen allergy patients that responded to allergen immunotherapy. We show *ITGA4* (CD49d) and *CD86* increased following allergy immunotherapy, in addition to *KLRG1* previously reported. CD49d and CD86 therefore also enabled tracking expansion of ILC2_10_ in a completely different context, supporting their broad utility in differentiating conventional ILC2s from ILC2_10_ in human studies. *Ex vivo* expanded ILC2_10_ from peripheral blood did not express KLRG1, which may indicate KLRGI is downregulated in *ex vivo* cultures, or that KLRG1 is differentially expressed by ILC2_10_ in different tissues or contexts. The combination of CD49d and CD86 distinguished IL-10^+^ ILC2s from conventional ILC2s in both studies, however, and our findings support these markers can be used to isolate them or track ILC2_10_ responses in human studies.

A wide range of prophylactic therapeutics have been explored to treat and prevent aGVHD, including post-transplant cyclophosphamide, calcineurin inhibitors like cyclosporine and other immunosuppressants like mycophenolate, anti-thymocyte globulin and sirolimus. While these treatments dramatically reduced incidence of aGVHD, development of aGVHD remains a major clinical challenge, affecting 30-50% of HSCT recipients(19–24). In a recent multicenter review which included a wide range of different treatment modalities, more than half of HSCT patients required a hospital stay within 100 days of transplant, with the primary reason being development of aGVHD, with severe GVHD (grades III-IV) occurring in 41.9% of patients(20, 21). Patients diagnosed with severe aGVHD were primarily managed with increased steroid doses (51.3% of patients), however, at follow-up, 52.8% of these patients were deceased(20, 21). Therefore, novel therapies are greatly needed to improve success of HSCT.

Here we examined direct effects of ILC2_10_ on allogeneic T cells that underlie GVHD pathology, which had not previously been explored. Adoptively transferred human ILC2_10_ suppressed allogeneic CD4^+^ Th1 and CD8^+^ Tc1 cells across multiple tissue compartments, and we noted an inverse relationship between ILC2s and CD4^+^Th1 cells in peripheral blood of HSCT recipients protected from GVHD. These findings collectively support ILC2_10_ have direct suppressive capacity that can be harnessed in cell therapies, and also that ILC2_10_ immunoregulatory activity extends beyond effects in the intestine. Prior mouse studies of conventional ILC2s reported ILC2-derived amphiregulin promotes intestinal repair and ILC2-mediated recruitment of MDSCs to the gut limits intestinal GVHD pathology locally, an important distinction with what we observed with human ILC2_10_ (27, 46). In support of ILC2_10_ having similar reparative functions to conventional ILC2s, however, expanded human ILC2_10_ expressed very high levels of amphiregulin.

While it would have been interesting to explore human ILC2_10_ effects on MDSCs or other immune populations, this is not possible using the xenogeneic GVHD model, which is almost exclusively engrafted by human T cells, with very limited contribution from myeloid cells, NK cells or B cells. Additional protective mechanism therefore likely exist, and could be explored in humanized models that support engraftment of other immune cells in future studies. However, use of this model enabled identification direct ability of ILC2_10_ to suppress allogeneic CD4^+^ and CD8^+^ T cells responses across multiple anatomical locations. Whether human ILC2_10_-mediated T cell regulation has broader cell therapy applications in T cell-driven pathologies, such as transplant rejection or T cell-driven autoimmune disorders will be an important area to explore moving forward.

Human ILC2_10_ suppress allogeneic xenoGVHD severity to similar levels as previously reported with polyclonal Tregs in this model(57). In clinical trials, evidence of Tregs limiting allo-immune responses was noted. However, use of polyclonal Tregs has not achieved significant response rates, with most patients still developing GVHD and requiring global immunosuppression(25, 26), highlighting the need for more effective therapies. We show that ILC2_10_ do not inhibit T cell engraftment, as was reported with Tregs in this model (48), but instead inhibit T cell proliferation, effector functions and tissue trafficking. Therefore, ILC2_10_-protective mechanisms are distinct from those observed with Treg cell therapy.

From a cell manufacturing perspective, ILC2_10_ might also have several advantages over Tregs. Tregs are typically sourced from the transplant donor or recipient, necessitating time sensitive isolation and expansion(58, 59). Our study using allogeneic ILC2_10,_ and a mouse study that demonstrated 3^rd^ party ILC2s can protect from intestinal damage(27, 46) support the possibility of using 3^rd^ party donors as an ‘off-the-shelf’ source of ILC2_10_, which would have advantages over current Treg approaches. Further, Tregs have limited expansion potential, whereas we show ILC2_10_ from peripheral blood exhibit robust expansion *ex vivo*. In addition to these cell manufacturing advantages, human ILC2_10_ have very high co-expression of amphiregulin, representing another potential benefit over human Tregs, which unlike mouse Tregs do not produce amphiregulin endogenously(27, 46). It will therefore be interesting to compare and contrast ILC2_10_ and Treg immunoregulatory mechanism in future studies, and explore whether dual regulatory and tissue-protective functions of human ILC2_10_ could offer an additional therapeutic advantage. What is clear however, is that ILC2_10_ display regulatory functions, and can limit inflammation in diverse contexts, therefore providing a strong rational to explore ILC2_10_-based cell therapies for a wide range of pathologies.

## Methods

### Human PBMC Isolation

Fresh blood was collected in EDTA collection tubes (BD Biosciences). In some instances, frozen PBMC samples were obtained from the Messner Allogeneic Transplant Program Biobank. PBMCs were isolated using Lymphoprep(StemCell Technologies) per manufacturer instructions.

### Flow Cytometry

Surface marker staining was performed for 30min after 15min blocking with human TruStain FcX(BioLegend). Cells were washed in FACS buffer(PBS+2% FCS) and fixed in 2%paraformaldehyde in PBS(ThermoFisherScientific). For intracellular staining, cells were fixed and permeabilized using FOXP3/Transcription Factor Staining set(eBioscience) and incubated with intracellular antibodies at room temperature(RT) for 30min. Samples were acquired on a LSR Fortessa(BD Biosciences) and data analyzed using FlowJo v10 software. Antibody details listed in **Supplemental Table 3**.

### Human ILC Sorting

PBMCs were stained with human TruStain FcX(BioLegend) and incubated with a cocktail of lineage antibodies conjugated to the FITC listed in **Supplemental Table 4**. Cells are washed in FACS buffer, resuspended in EasySep Buffer(StemCell Technologies) and enriched using the EasySep FITC Positive Selection Kit II(StemCell Technologies) per manufacturer instructions. Enriched cells were stained with antibody cocktail(**Supplemental Table 5**), and sorted using a FACSAria Fusion(BD Biosciences).

### Human ILC Expansion

Sorted human ILC2s and ILC3s were cultured in complete X-VIVO 15 media(Lonza) supplemented with 5% human AB serum(Sigma), 100U/mL Penicillin-Streptomycin(Gibco) and 1x GlutaMAX(Gibco).

ILC2s were expanded using 100 IU/mL IL-2(SteriMax), 20ng/mL IL-7 and 20 ng/mL IL-33(BioLegend).

ILC3s were expanded using 100 IU/mL IL-2, 20ng/mL of IL-1β, IL-7 and IL-23(BioLegend). CD56^bright^ and CD56^dim^ NK cells were expanded in NK MACS media(Miltenyi) supplemented with 5% human AB 500 IU/mL of IL-2, and 20ng/mL of IL-15 and IL-18(Biolegend).

### Cytokine and Chemokine Assays

For intracellular cytokine analysis, 2×10^5^ cells were stimulated with Cell Stimulation Cocktail(eBioscience) for 6hrs, with GolgiStop and GolgiPlug(BD Biosciences) added after 2hrs. Cells were then intracellularly stained with antibodies as detailed in **Supplemental Table 6**. For secreted factor analysis, 1×10^5^ cells were plated in complete X-Vivo media and stimulated with 100 IU/mL of IL-2. After 16hrs, supernatant was collected and stored at -80°C. Analytes were measured using the 12-plex LegendPlex Human Th Cytokine Panel(BioLegend) or the 13-plex LegendPlex Human Proinflammatory Chemokine Panel 1(BioLegend) on thawed supernatant samples per manufacturer’s instructions.

### Xenogeneic GVHD Mouse Model

Six-to-ten-week-old NOD.Cg-*Prkdc^scid^Il2rg^tm1Wjl^*/SzJ(NSG) mice were given 150cGy irradiation one day prior to intravenous tail vein injection of 1×10^7^ freshly isolated PBMCs with or without 1×10^7^ expanded allogeneic ILC2s in PBS at the same time or with ILC2s administered 9 days later (upon GVHD onset) (47, 60). Control mice were injected with PBS either at Day 0 or Day 9. Mice were monitored daily for symptoms of GVHD including weight loss, fur loss and skin inflammation, hunch, activity, and pain, scored on a scale of 0–3. At endpoint, blood, bone marrow and spleens were harvested as previously described(60, 61). For intestinal immune cells, intestines were flushed, sliced, and rinsed in 1xPBS. Tissue was cut into segments, added to 2mM EDTA in 1xPBS and incubated for 30min at RT. Tissue was strained and the remaining tissue was added to digestion buffer containing 1,650NPA-U BP protease(VitaCyte), 2,500CDA-U CollagenaseMA(VitaCyte), and 100µg/mL DNaseI(StemCell Technologies) in Hank’s Balanced Salt Solution with calcium(Wisent Bio Products). Tissue was then cut and incubated at 37°C for an additional 30min. Tissue was passed through a 70µm filter and washed in RPMI (Gibco)+5%FCS twice.

For graft-versus leukemia experiments, six-to-ten-week-old NSG mice were given 250cGy irradiation one day prior to intravenous tail vein injection of 2×10^6^ MV4-11 cells per mouse. Five days later, mice were given 5×10^6^ PBMCs alone or with 5×10^6^ expanded human ILC2s. Mice were sacrificed 14 days after PBMC injection, and the presence of MV4-11 cells in bone marrow was assessed.

### Histology

Harvested spleen, lung and colon tissue were fixed for 3 days in 10% neutral buffered formalin and stored in 70% ethanol. Paraffin embedding, tissue slicing, H&E and immunohistochemical staining was performed by the UHN Pathology Research Program Laboratory. Quantification of cells expressing IHC markers was performed using the HALO Image Analysis Platform and reported as a percent of all cells.

### Co-Cultures

Naïve CD4^+^ and total CD8^+^ T cells were isolated from the healthy donor blood using the EasySep Human Naïve CD4^+^ T cell or the EasySep Human CD8^+^ T cell isolation kit(StemCell Technologies). T cells were plated at 5×10^4^ per well and stimulated with human CD3/CD28 T-Activator DynaBeads(Gibco) at 1:8 beads:T cell ratio. Expanded ILC2s were added 1:1 to the T cells. T cells were assessed for markers after 72hrs by flow cytometry. For ILC supernatant experiments, expanded ILC2s were plated 5×10^4^ cells/mL in complete X-Vivo media without cytokines. After 16hrs, supernatant was harvested, centrifuged at 1500rpm for 10min and stored at -80°C. Supernatant was thawed on day of T cell cultures, and added 1:1 with T cells. Fresh supernatant was added every 24hrs for 3days. For inhibitor experiments, UltraLeaf anti-human IL-4, IL-9, IL-10, and IL-13(BioLegend) were added at 10µg/mL. ARL67156(Tocris) and PBS12379(Tocris) were added at 1µg/mL to inhibit activity of CD39 and CD73, respectively. Inhibitors were added at day zero and every 24hrs for 72hrs.

### CITE-seq on expanded ILCs and analysis

1×10^6^ expanded CD56^bright^, CD56^dim^, ILC2s and ILC3s were washed twice in 1x Cell Staining Buffer(BioLegend) and incubated in TruStain FcX for 15min. Cells were incubated with a TotalSeq-C antibody cocktail as well as hashtag antibodies to allow for post sequencing differentiation of distinct expanded ILC subsets(BioLegend, **Supplemental Table 1**) for 30min. Antibody concentrations and background signal were determined as previously described(62). Cells were washed 3x in Cell Staining Buffer and resuspended in 1xPBS with 0.04% BSA(Millipore Sigma). Samples were prepared for sequencing using the 10X Genomics Single Cell 5’ v2 platform in accordance with manufacturer’s instructions for capture of 12,000 cells per samples. Reverse transcription, cDNA amplification and sequencing libraries using the 10X Genomics Single Cell 5’ v2 reagents. Samples were sequenced to a depth of 40,000 reads. Read alignment to the reference human genome (GRCh38/hg38) and gene expression matrices were generated using CellRanger v6.1.2. Single cells were filtered to exclude cells that expressed >10% mitochondrial content, <1000 total transcripts and <200 unique genes. Data was log normalized, principal component analysis was preformed, and cells were clustered by the top 20 principal components with the Louvain community algorithm using Seurat’s FindNeighbors and FindClusters(63). Cell cycle genes were assigned using Seurat’s CellCycleScoring and regressed out during scaling. Clusters were visualized using the Uniform Manifold Approximation and Projection(UMAP)(64). Clusters were assigned to ILC subsets based on expression of hashtag antibodies, lack of lineage markers and expression of subset associated markers(**Figure 6a**). Cluster defining markers were identified using FindMarkers in Seurat. Differential expression of genes was visualized using EnhancedVolcano(65).

## Data availability

CITEseq dataset of expanded human ILCs will be made publicly available via NCBI GEO at the time of publication. Publicly available single-cell RNA-sequencing dataset of ILCs in patients undergoing grass-pollen allergy immunotherapy was originally published as part of studies by Golebski *et al* (13).

## Statistics

Statistical significance was determined by Kruskal-Wallis test or one-tailed Mann-Whitney test. The log-rank(Mantel-Cox) test was used for Kaplan-Meir curves. Analysis of correlational data was calculated using Spearman correlation. The number of replicates is represented by *n* and is indicated in each figure legend. *p<0.05;**P<0.01;***P<0.001;****P<0.0001;*ns,* not significant. Error bars represent standard deviation unless otherwise stated. Data analysis was preformed using GraphPad Prism v9.

## Study Approval

Patient study protocols were approved by the Research Ethics Board at University Health Network in accordance with the Helsinki Declaration (UHN REB 19-6351), with all patients provided written, informed consent. Healthy peripheral blood was obtained from donors through the Canadian Blood Services Blood4Research program, with each donor providing written, informed consent(UHN REB 17-6229, CBS Approved Study 2020-047). Animal study protocols were approved by the University Health Network Animal Care Committee(UHN AUP 6203) and all ethical regulations were followed.

## List of Supplementary Material

**Supplemental Figure 1.**
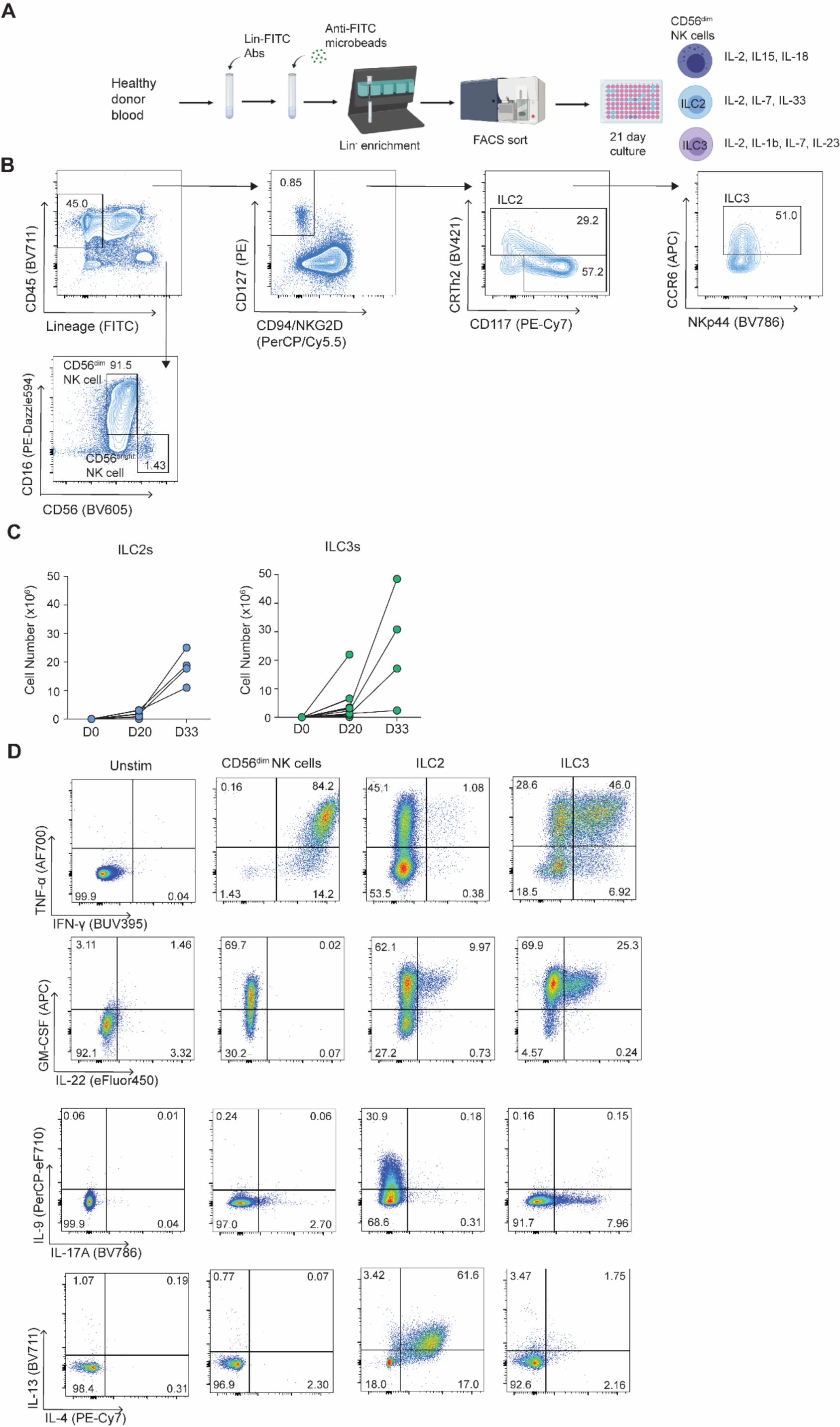
Flow cytometry gating strategy for ILC sorting. To explore the role of human ILC2s in GVHD, human ILCs had to be isolated from peripheral blood using flow cytometry and expanded ex vivo using ILC subset-specific cytokines as they are present in very low abundance in peripheral blood. (A) Overview of isolation and expansion of human ILC subsets. Briefly, total PBMCs are isolated from healthy donor blood. PBMCs are incubated with FITC-conjugated linage antibodies. Lineage antibodies added are CD3 (OKT3), CD3 (UCHT1), CD4, CD8a, CD14, CD15, CD19, CD20, TCRꭤꞵ, TCR𝛾𝛿, CD33, CD34, CD203c, FCεRI, CD79a and CD138. Anti-FITC magnetic microbeads are added, and cells are enriched for lineage^-^ cells using StemCell magnets. Enriched cells are FACS sorted for CD56^dim^ NK cells, ILC2s and ILC3s. ILC2s are cultured with 100U/mL IL-2, 10ng/mL IL-7 and 10ng/mL IL-33, ILC3s are cultured with 100U/mL IL-2, 10ng/mL IL-7, 10ng/mL IL-1ꞵ, and 10ng/mL of IL-23 and CD56^dim^ NK cells are cultured with 500U/mL IL-2, 10ng/mL IL-15, and 10ng/mL of IL-18. (**B**) Representative gating strategy for ILC subsets. All ILCs are gated as Live, CD45^+^ and Lin^-^. NK cells are then gating as CD56^+^ CD16^+^ for CD56^dim^ NK cells and CD56^bright^ CD16^-^ for CD56^bright^ NK cells. Helper ILCs are then sorted as live, lineage^-^ CD94^-^NKG2D^-^CD127^+^ with ILC2s being CRTh2^+^ CCR6^-^ and ILC3s being CRTh2^-^CD117^+^CCR6^+^. (**C**) Cell expansion yields of ILC2s and ILC3s at day 20 and day 33. (**D**) Expanded ILCs intracellular cytokine staining with unstimulated ILCs as the gating control.

**Supplemental Figure 2.**
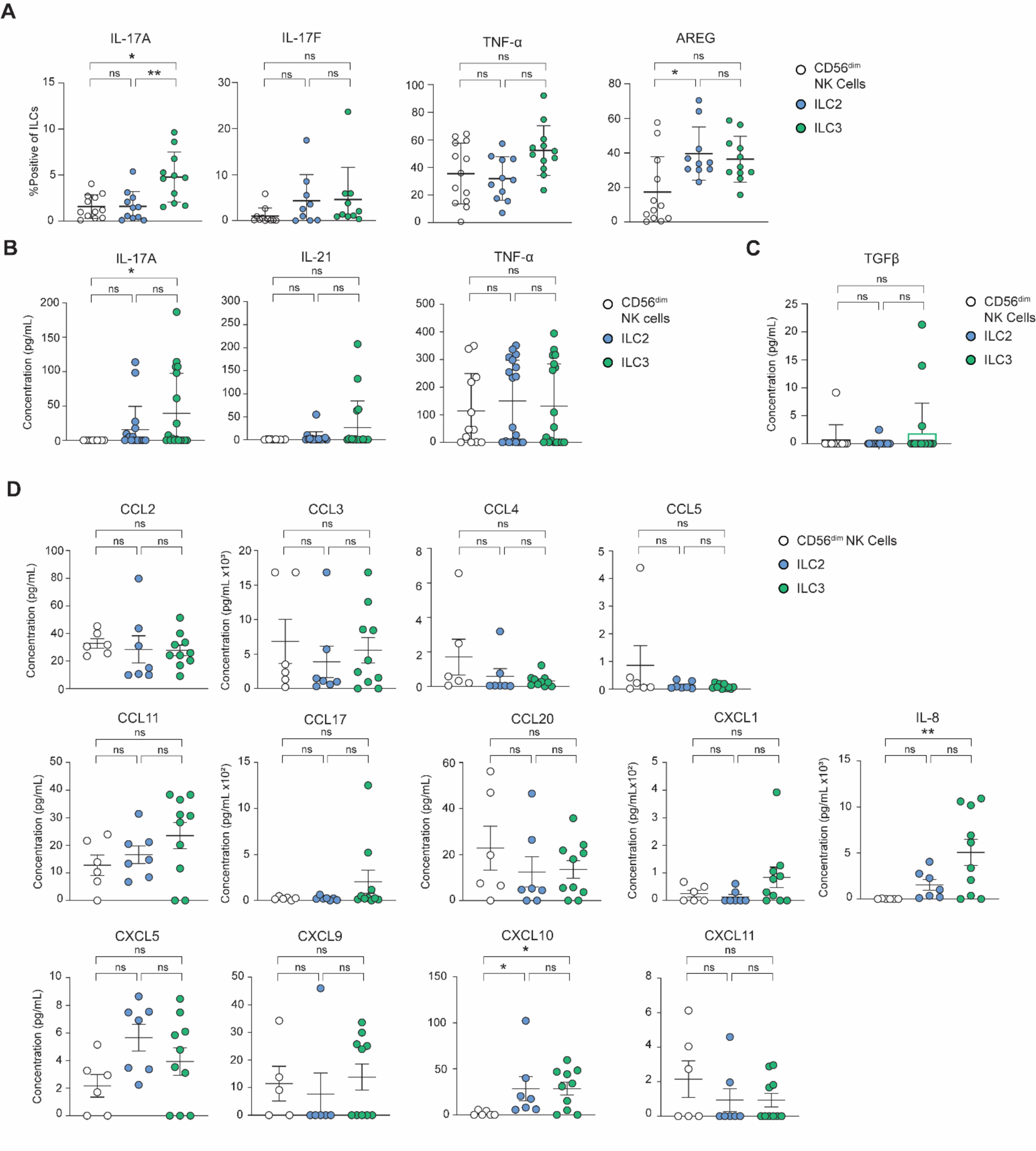
Analysis of chemokines and cytokines produced by expanded ILC populations. Further chemokine and cytokine analysis of ILC2s with an ILC2_10_ phenotype, as compared to NK cells and ILC3s. (**A**) Intracellular cytokine staining for IL-17A, IL-17F, TNF-α and Amphiregulin after 6-hour stimulation with phorbol 12-myristate 13-acetate and ionomycin and blocked with monensin and brefeldin A (n=13, 11 and 12 for CD56^dim^ NK cells, ILC2s and ILC3s respectively). (**B**) Cytometric bead array (CBA) analysis of secreted IL-17A, IL-21, and TNF-α after plating ILCs at a concentration of 2×10^5^ cells/mL for 16-hour stimulation with 100U/mL of IL-2 (n=13, 18 and 18 for CD56^dim^ NK cells, ILC2s and ILC3s respectively). (**C**) CBA analysis of secreted active TGF-β1 after plating ILCs at a concentration of 2×10^5^ cells/mL for 16-hour stimulation with 100U/mL of IL-2 (n=13, 18 and 18 for CD56^dim^ NK cells, ILC2s and ILC3s respectively). (**D**) CBA analysis of secreted chemokines after plating ILCs at a concentration of 2×10^5^ cells/mL for 16-hour stimulation with 100U/mL of IL-2. (n=6, 7 and 10 for CD56^dim^ NK cells, ILC2s and ILC3s respectively).

**Supplemental Figure 3.**
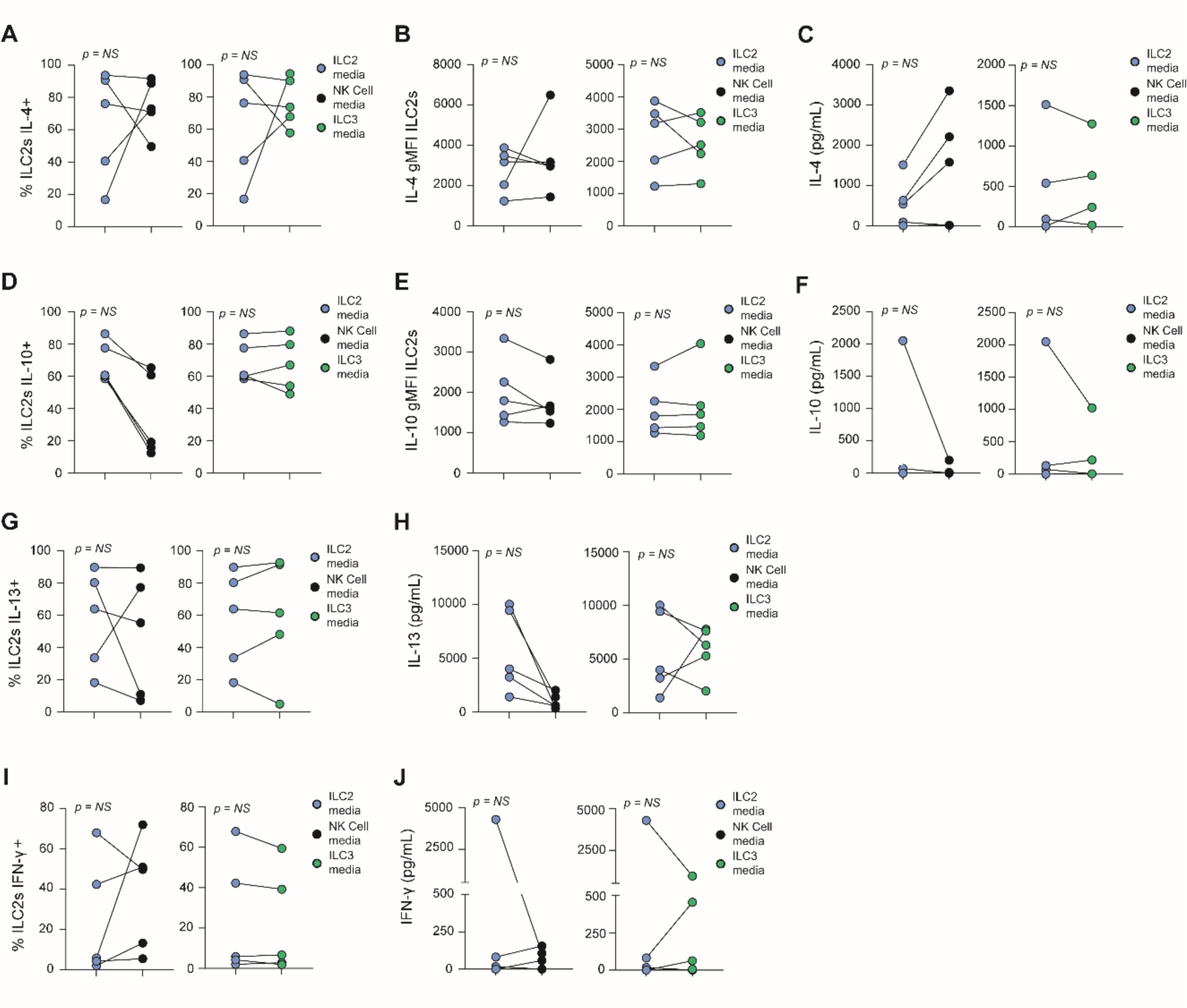
Stability of signature cytokines on ILC2s with ILC2_10_ phenotype upon culture in different cytokine conditions. ILC2_10_ were expanded in ILC2 medium (X-VIVO15 Complete with IL-2, IL-7, and IL-33). These ILC2_10_ were then removed from culture, washed, and plated in NK cell media (NK MACS medium with IL-2, IL-15, and IL-18) or ILC3 medium (X-VIVO15 Complete with IL-2, IL-7, IL-1β and IL-23). Intracellular cytokines were measured after 6-hour stimulation with phorbol 12-myristate 13-acetate and ionomycin and blocked with monensin and brefeldin A. **A,B** Expression of IL-4 on expanded ILC2_10_ cultured in NK cell medium or ILC3 medium compared to ILC2 medium. This was measured as the percent of ILC2_10_ positive for IL-4 (**A**) or the gMFI of IL-4 (**B**). (**C**) Cytometric bead array (CBA) analysis of changes in secreted IL-4 from ILC2s cultured in NK cell or ILC3 medium. **D,E** Expression of IL-10 on expanded ILC2_10_ cultured in NK cell or ILC3 medium compared to ILC2 medium measured as the percent of ILC2s positive (**D**) or gMFI (**E**) of IL-10. (**F**) CBA analysis of changes in secreted IL-10 on expanded ILC2s subsequently cultured in NK cell or ILC3 medium. **G,H** Expression of IL-13 on expanded ILC2_10_ cultured in NK cell or ILC3 medium compared to ILC2 medium measured as the percent of ILC2s positive (**G**) or gMFI (**H**) of IL-10. (**I**) Expression of IFN-γ on ILC2_10_ cultured in ILC2 medium by percent positive following stimulation compared to culturing in NK cell or ILC3 medium. (**J**) CBA analysis of changes in secreted IFN-γ from ILC2s cultured in NK cell or ILC3 medium.

**Supplemental Figure 4.**
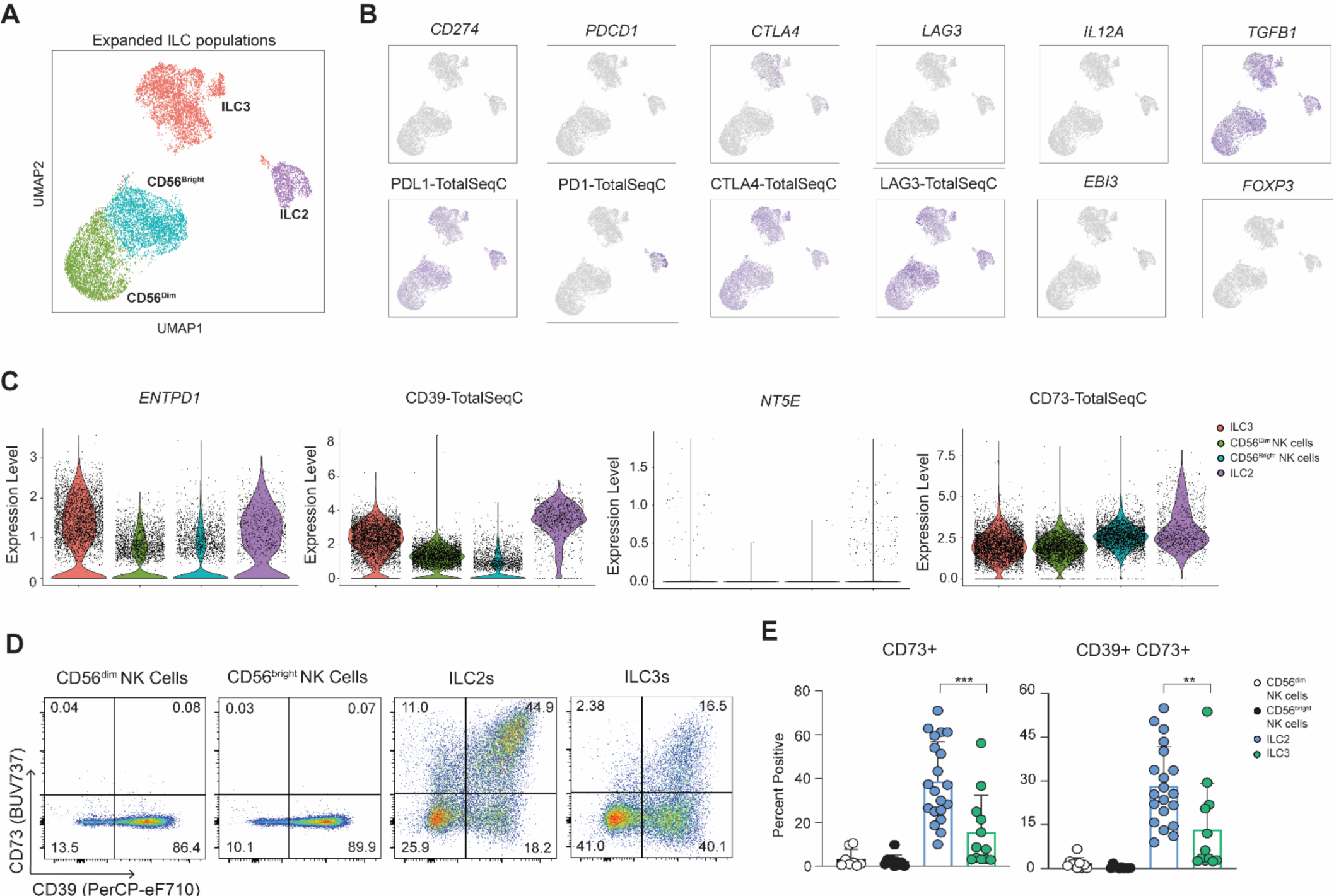
Expression of nTreg and Tr1 markers on expanded ILC2s with ILC2_10_ phenotype. (**A**) UMAP plots of expanded ILC subsets after CITE-seq. ILC subsets were isolated using FACS, expanded and then separately stained with an antibody cocktail. Each cell population was stained with a unique hashtag antibody to allow identification post sequencing. (**B**) Feature plots showing expression of markers linked to Treg and Tr1 cell function. Protein level expression is shown of PD-L1, PD-1, CTLA-4 and LAG-3 and RNA level expression of *CD274, PDCD1, CTLA4, LAG3, IL12A, EBI3, FOXP3* and *TGFB1* on expanded ILC subsets. (**C**) Violin plots of single cell expression of protein level CD39 and CD73 and RNA expression of *ENTPD1* and *NT5E* after expansion. Flow cytometry expression of CD39 and CD73 on expanded ILC subsets shown as representative flow plots (**D**) and summary graphs **(E)** (n = 10-20).

**Supplemental Figure 5.**
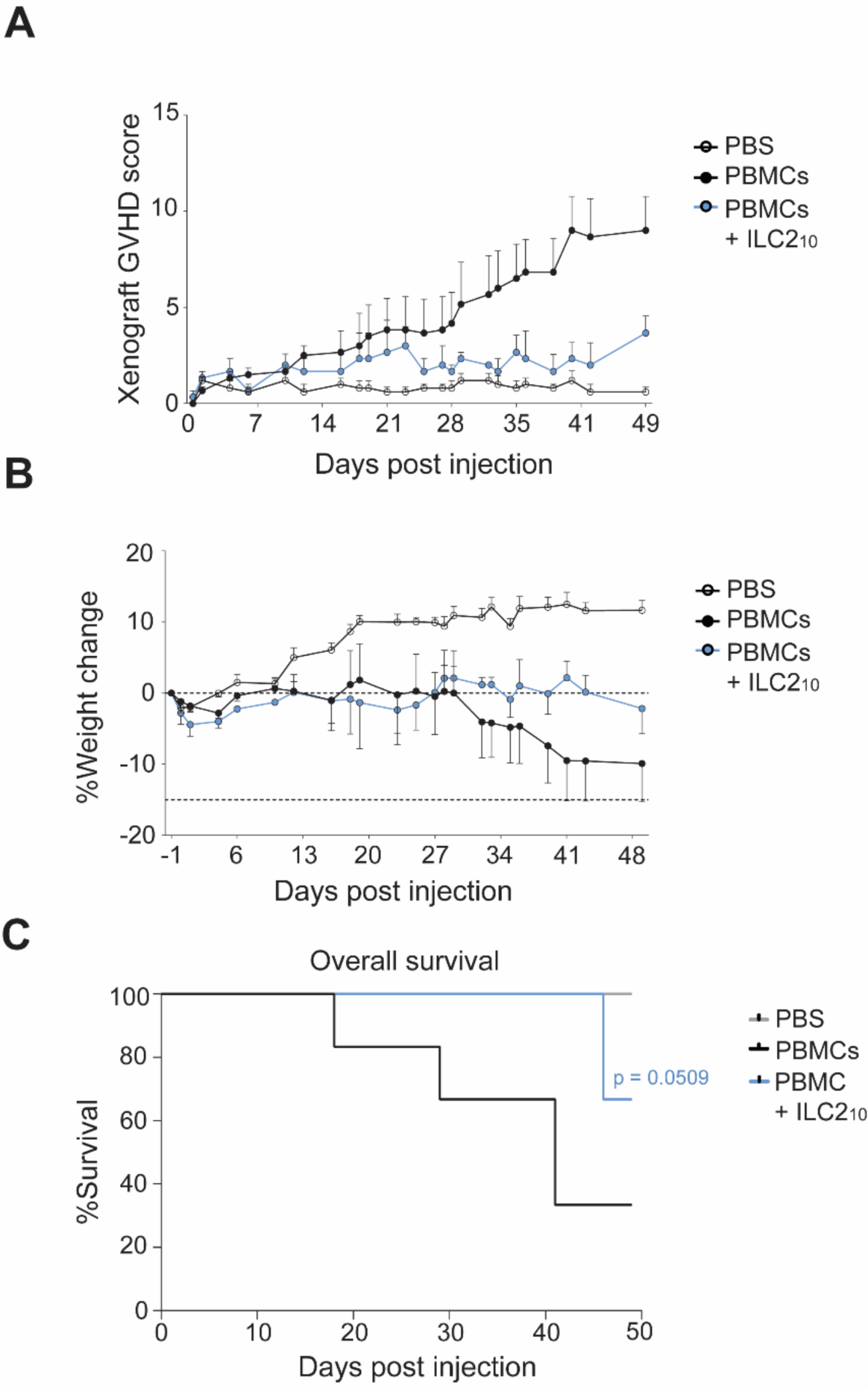
Expanded human ILC2_10_ suppress symptoms and improve survival of xenograft GVHD in male mice. (**A-C**) Xenogeneic GVHD score (**A**) weight loss (**B**) and overall survival **(C)** of male mice given PBS, 1×10^7^ human PBMCs to induce xenograft GVHD or 1×10^7^ PBMCs with an equal number of expanded human ILC2_10_ over 7 weeks (n=5).

**Supplemental Figure 6.**
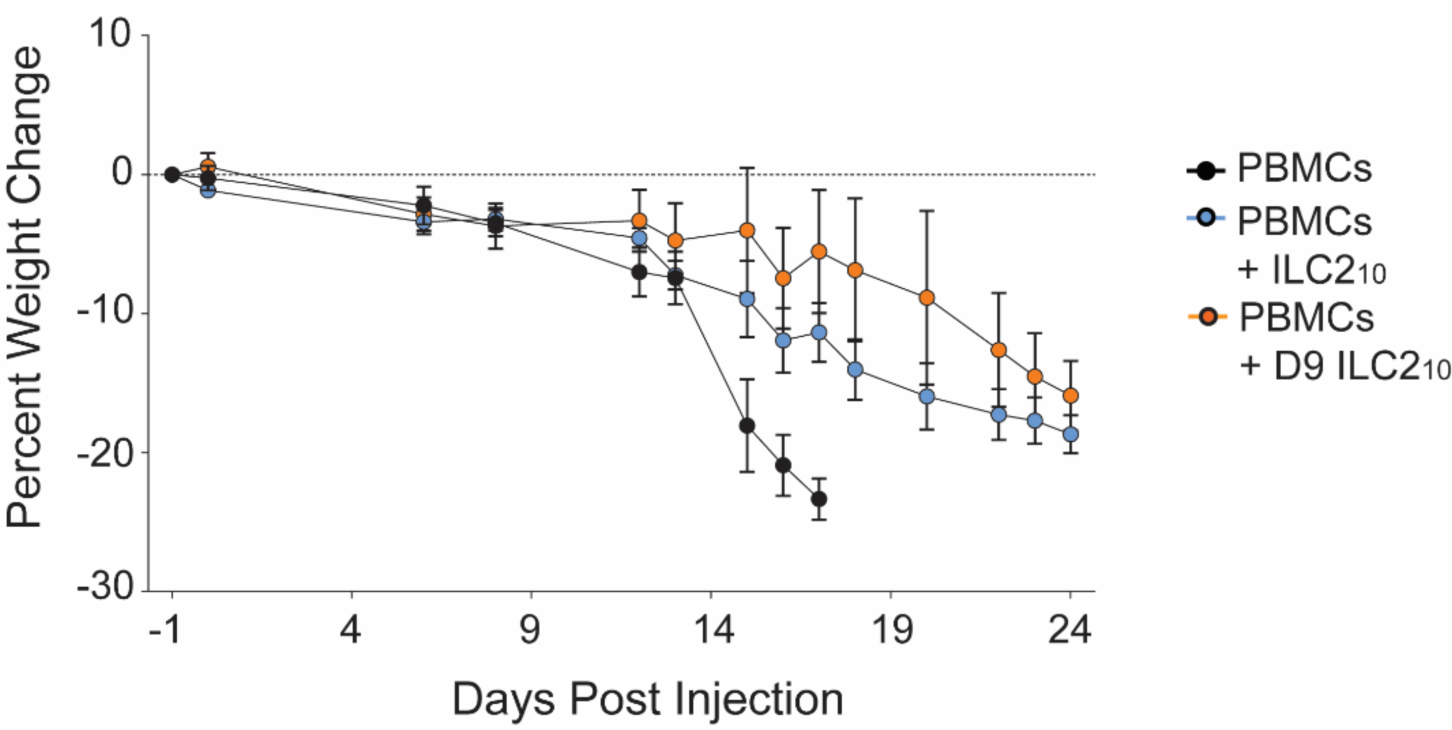
Delayed injection of expanded human ILC2_10_ improve weight-loss in xenoGVHD mice. Weight-loss over time of xenogeneic GVHD mice treated with expanded human allogeneic ILC2_10_ at day 0 or 9 days following PBMC injection (n=3 for PBMC+ILC2_10_ groups and n=5 for PBMCs alone).

**Supplemental Figure 7.**
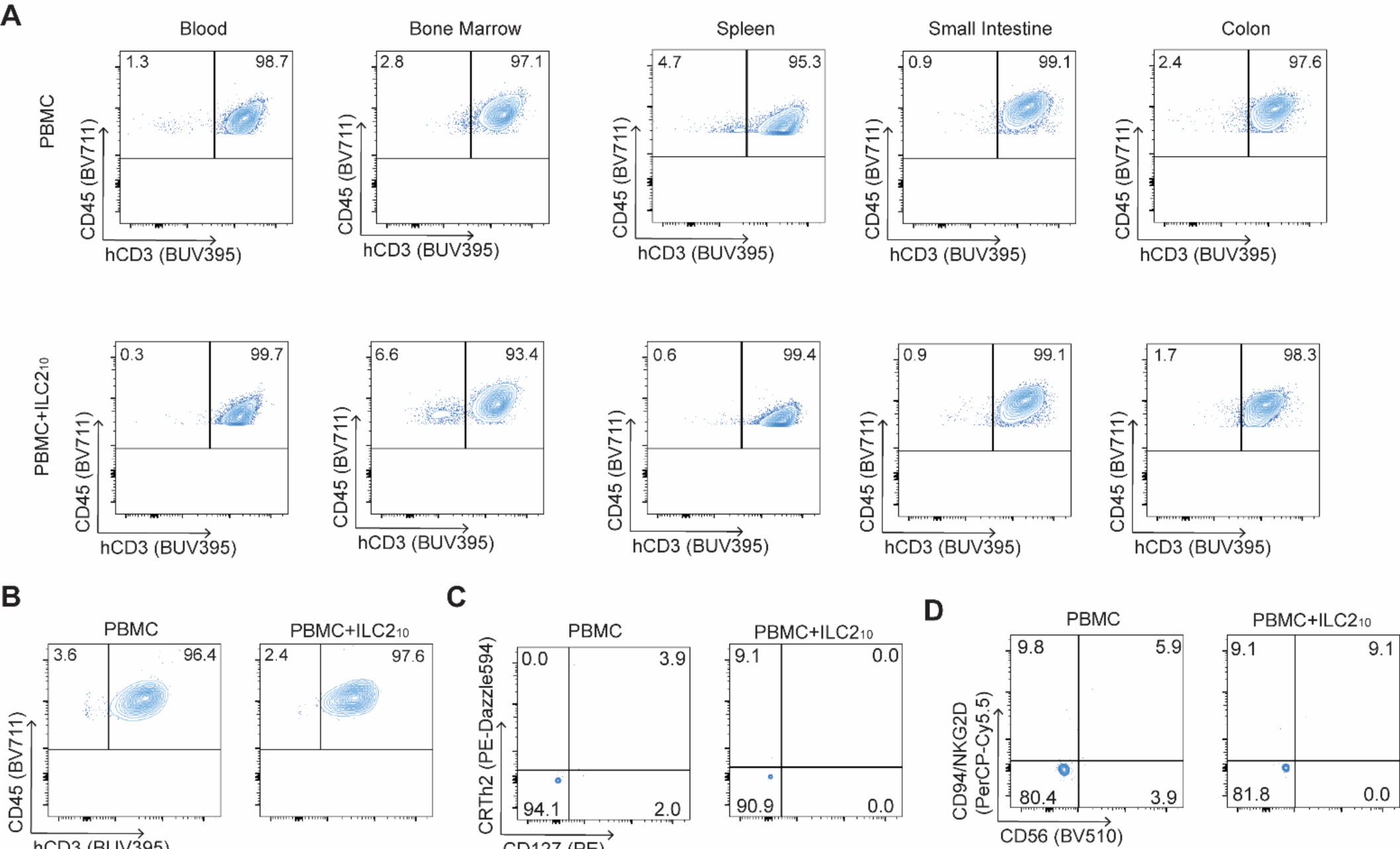
Engrafted human CD45^+^ cells are primarily T cells in xenogeneic GVHD mice. Flow cytometry plots of engrafted CD45^+^ human cells in NSG mice given PBMCs alone or with allogeneic expanded ILC2_10_. **A,B** Engraftment of human CD3^+^ T cells gated on CD45^+^ at endpoint (**A**) or day 14 post PBMC injection (**B**). **C,D** Proportion of human helper ILCs and ILC2s gated on CD45^+^CD3^-^ cells (**C**) and NK cells gated on CD45^+^CD3^-^ cells (**D**).

**Supplemental Figure 8.**
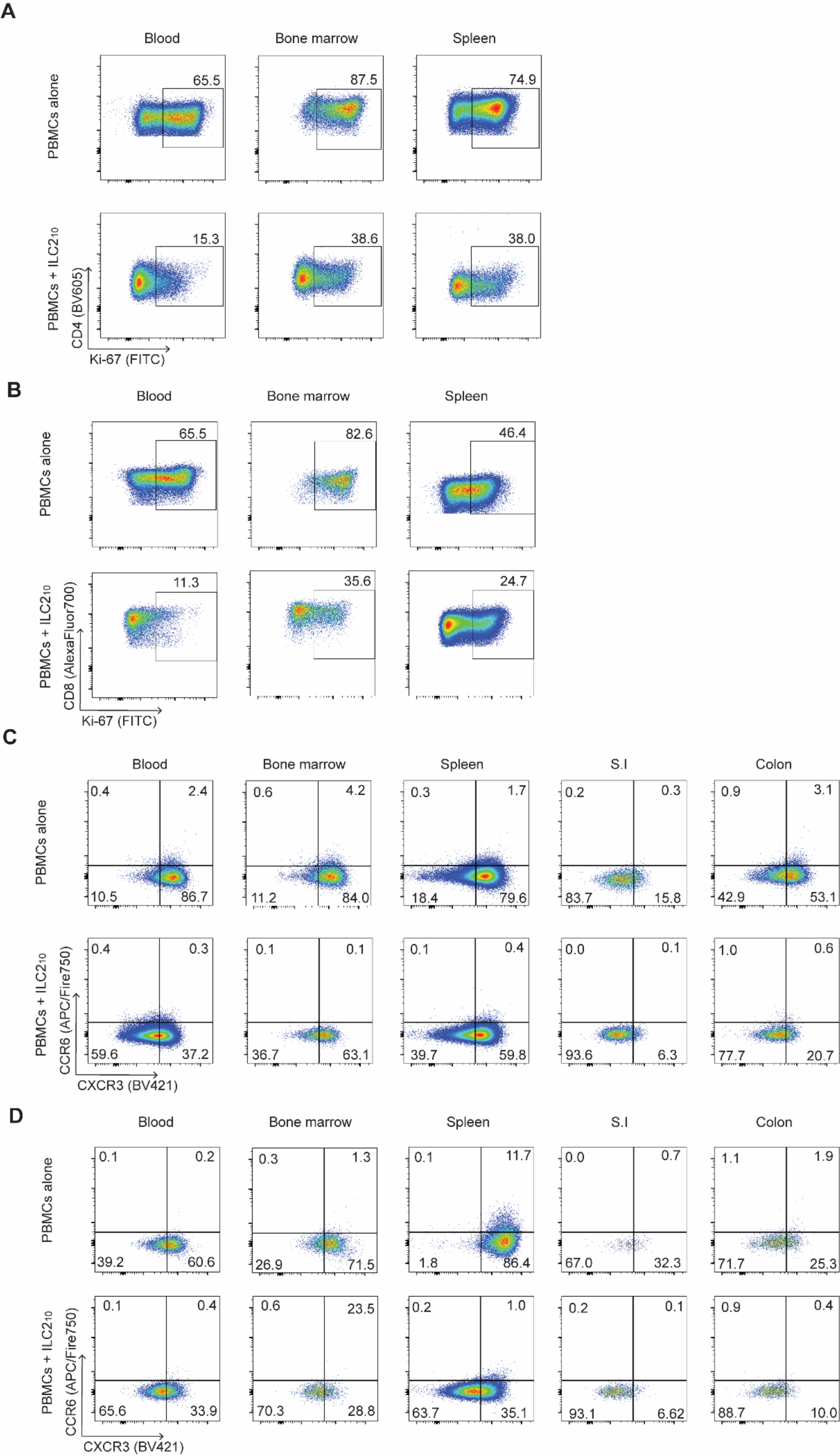
Representative flow cytometry analysis of Ki-67 and CXCR3 on CD4^+^ and CD8^+^ T cells *in vivo* across tissues. A,B. Representative FACS plots showing Ki-67 expression on CD4^+^ (**A**) and CD8^+^ (**B**) T cells in the blood, bone marrow and spleens of NSG mice given PBMCs or PBMCs with ILC2_10_ infusion. **C,D** Representative FACS plots showing CXCR3 expression on CD4^+^ (**C**) and CD8^+^ (**D**) T cells in the blood, bone marrow, spleen, small intestines (S.I.) and colon of NSG mice given PBMCs or PBMCs with ILC2_10_ infusion.

**Supplemental Figure 9.**
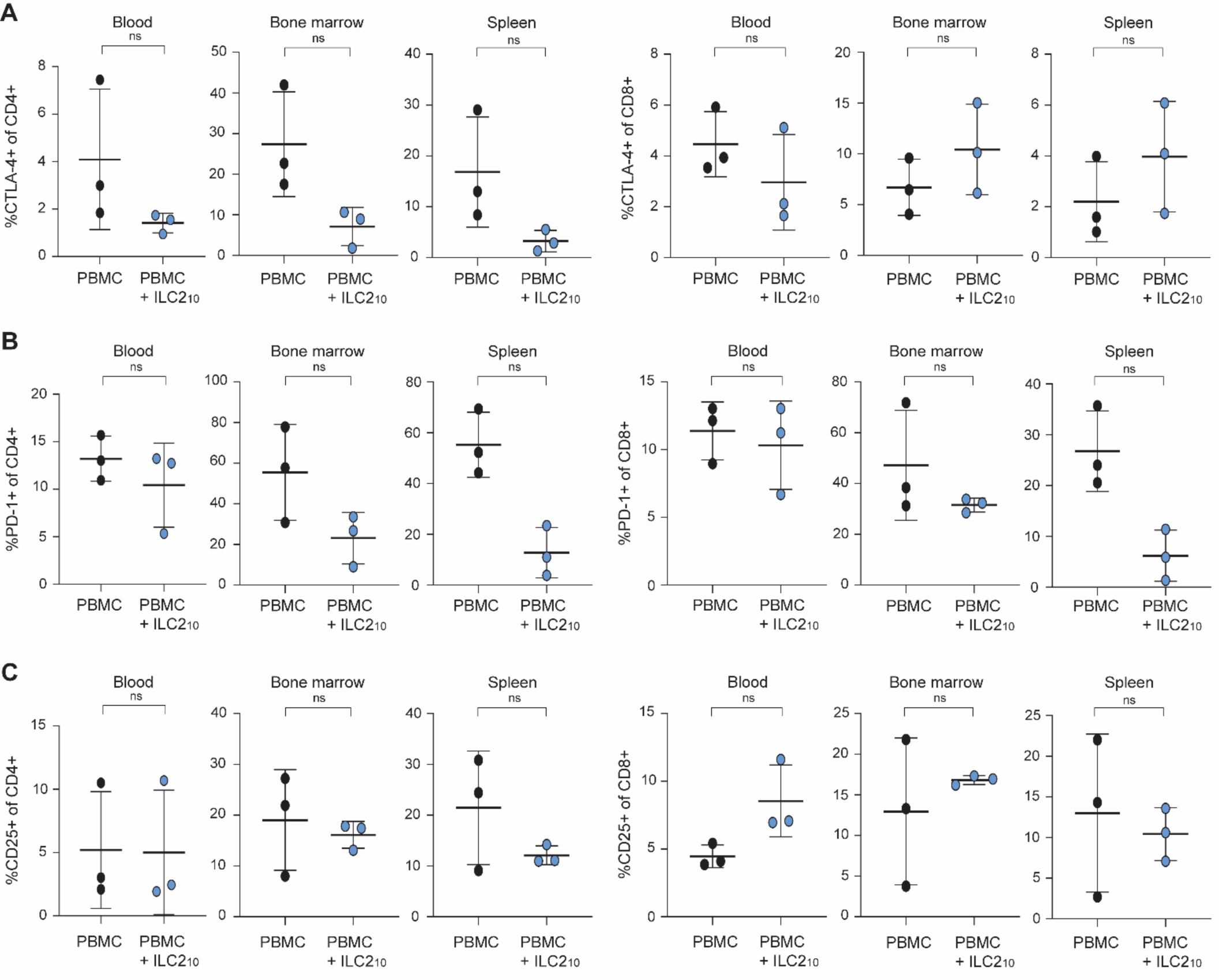
Activation and checkpoint molecule expression on CD4^+^ and CD8^+^ T cells. **A-C** Expression of CTLA-4 (**A**) PD-1 (**B**) and CD25 (**C**) on CD4^+^ and CD8^+^ T cells in the blood, bone marrow and spleens of NSG mice given PBMCs or PBMCs with ILC2_10_ infusion (n = 3).

**Supplemental Figure 10.**
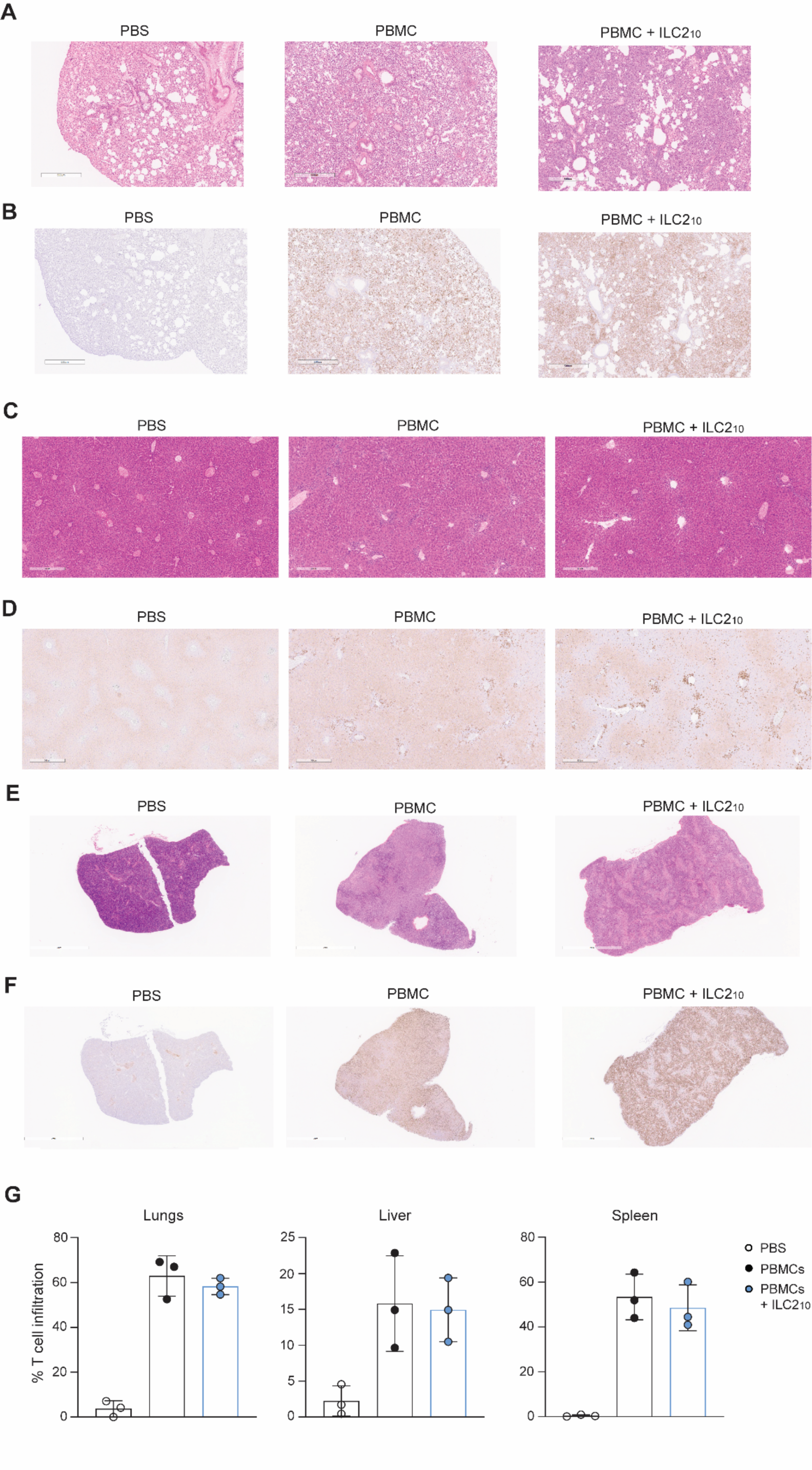
H&E Staining and human CD3 immunohistochemical staining tissues of xenoGVHD mouse model. **A,B** Representative H&E (**A**) and human CD3 (**B**) staining on lungs from NSG mice given PBS, PBMCs or PBMCs with ILC2_10_ taken at endpoint 8x magnification. **C,D** Representative H&E (**C**) and human CD3 (**D**) staining on livers from NSG mice given PBS, PBMCs or PBMCs with ILC2_10_ taken at endpoint at 8x magnification. **E,F** Representative H&E (**E**) and human CD3 (**F**) staining on spleens from NSG mice given PBS, PBMCs or PBMCs with ILC2_10_ taken at endpoint 2x magnification. (**G**) Average T cell infiltration in the lungs, liver and spleen of mice treated with PBS, PBMCs or PBMCs + ILC2_10_. Quantified using CD3 immunohistochemistry and the HALO algorithm (n=3).

**Supplemental Figure 11.**
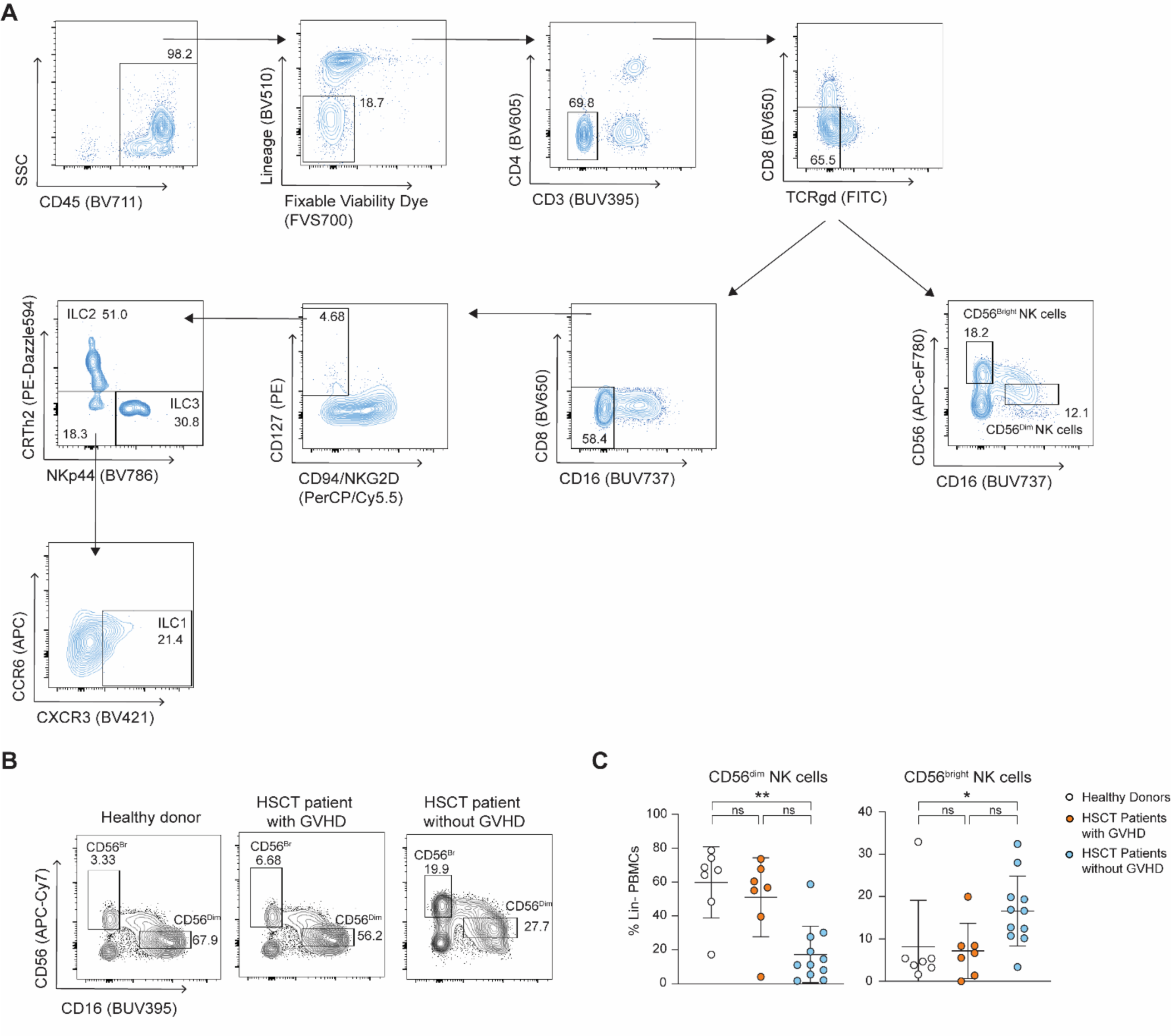
ILC Gating strategy to analyze ILCs in HSCT recipient peripheral blood. PBMCs from healthy donors, and HSCT patients with or without aGVHD were assessed for ILC proportions and phenotypes using flow cytometry. (**A**) Lineage markers are CD14, CD19, CD20, CD33, CD34, CD123, CD138, CD303, and FCεRI. Cells are gated as CD45^+^, Lineage^-^, Live, CD4^-^, CD3^-^, CD8^-^, and TCR𝛾𝛿^-^. NK cells are then gating as CD56^+^ CD16^+^ for CD56^dim^ NK cells and CD56^bright^ CD16^-^ for CD56^bright^ NK cells. Helper ILCs are then gated as CD16^-^CD94^-^NKG2D^-^CD127^+^ with ILC1s gated as CD117^-^CRTh2^-^CXCR3^+^, ILC2s as CRTh2^+^ and ILC3s as CRTh2^-^CD117^+^. **B,C** Representative flow plots (**B**) and summary graphs (**C**) of CD56^dim^ CD16^+^ and CD56^bright^ CD16^-^ NK cells in PBMCs, as a proportion of live CD45^+^ lineage negative cells.

**Supplemental Figure 12.**
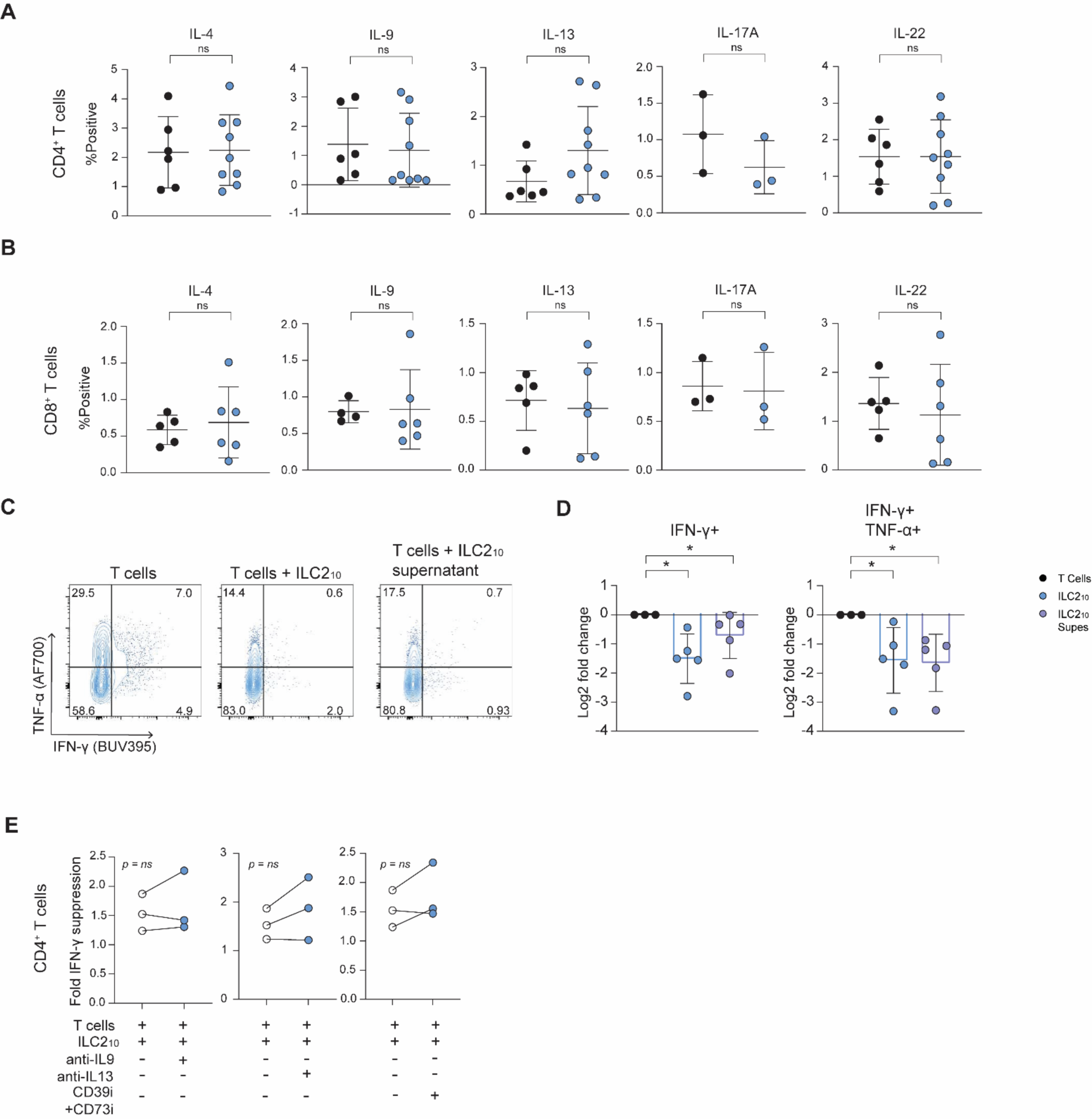
Effect of ILC2_10_ supernatant and addition of various inhibitors to ILC2_10_-T cell co-cultures. **A,B** Summary graphs of intracellular IL-4, IL-9, IL-13, IL-17A and IL-22 on CD4^+^ (n=6 T cells, n=9 ILC2_10_)(**A**) and CD8^+^ (n=5 T cells, n=6 ILC2_10_) (**B**) T cells after co-culture with ILC2_10_. **c,d** Representative FACS plots (**C**) and summary graphs (**D**) of IFN-γ expression by CD4^+^ T cells cultured alone, with ILC2_10_ or with ILC2_10_ supernatant (n=5). (**E**) Changes in fold IFN-γ expression after co-culture of CD4^+^ T cells with ILC2_10_ with the addition of IL-9 or IL-13 antibodies or with CD39 and CD73 inhibitors on CD4^+^ T cells (n=3).

**Supplemental Figure 13.**
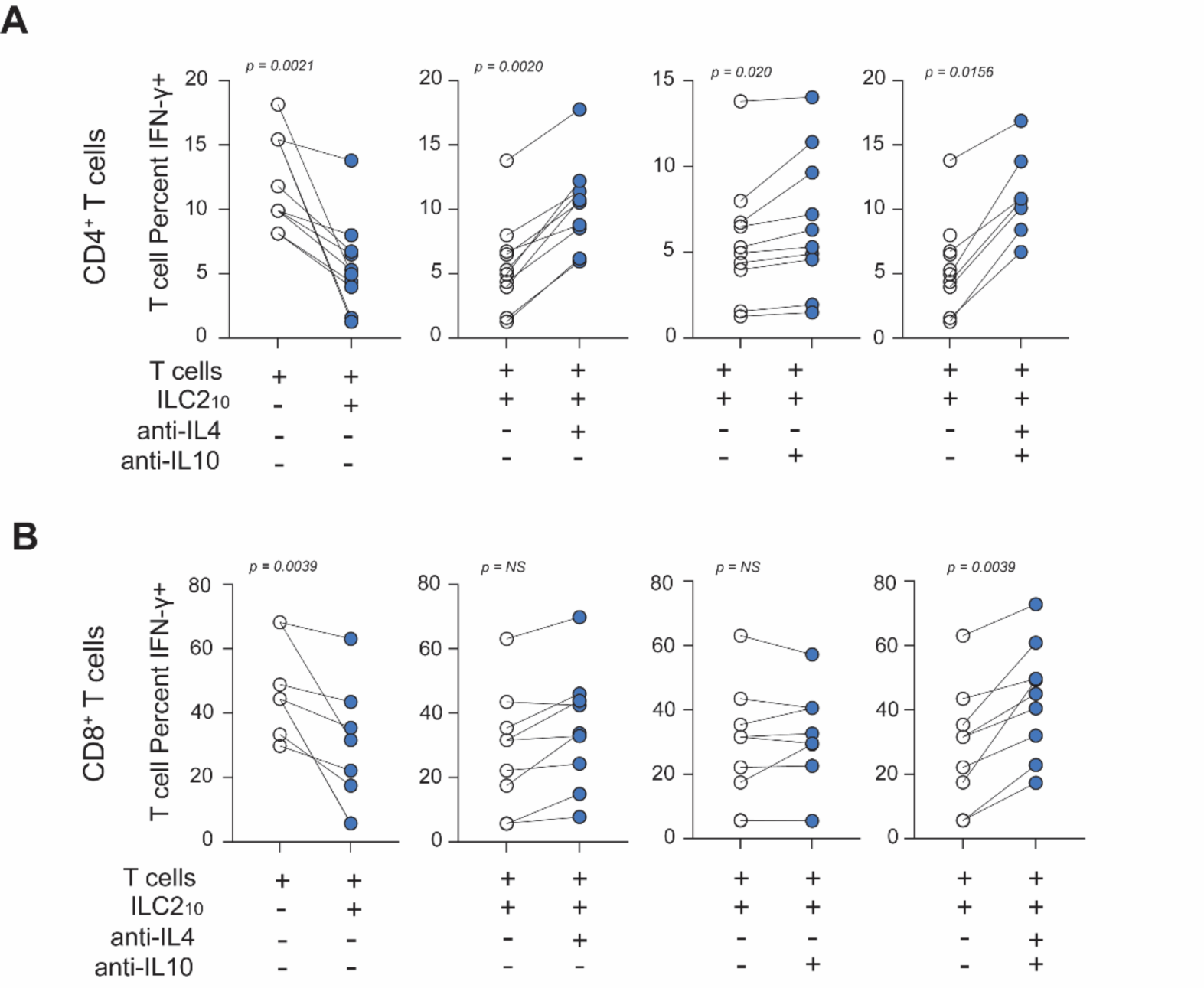
Non-normalized IFN-γ expression on CD4^+^ and CD8^+^ T cells cultured with ILC2_10_. (**A**) Percent expression of IFN-γ by CD4^+^ T cells cultured alone or with ILC2_10_ in the presence or absence of neutralizing antibodies towards IL-4, IL-10 and IL-4 and IL-10 together (n=10 from 5 independent experiments**) (B**) Percent expression of IFN-γ by CD8^+^ T cells cultured alone or with ILC2_10_ in the presence or absence of neutralizing antibodies towards IL-4, IL-10 and IL-4 and IL-10 together (n=7 from 5 independent experiments)

**Supplemental Table 1:**
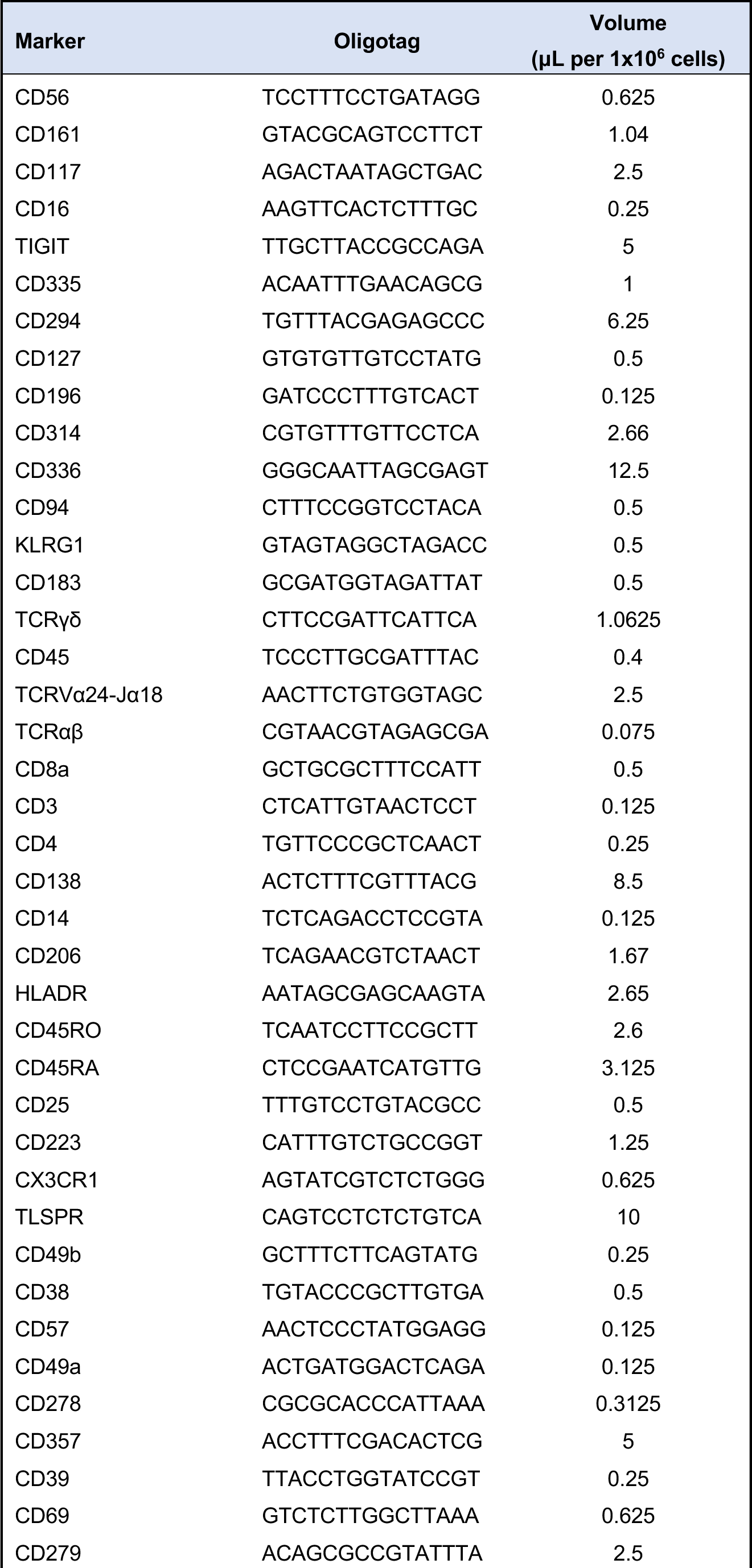
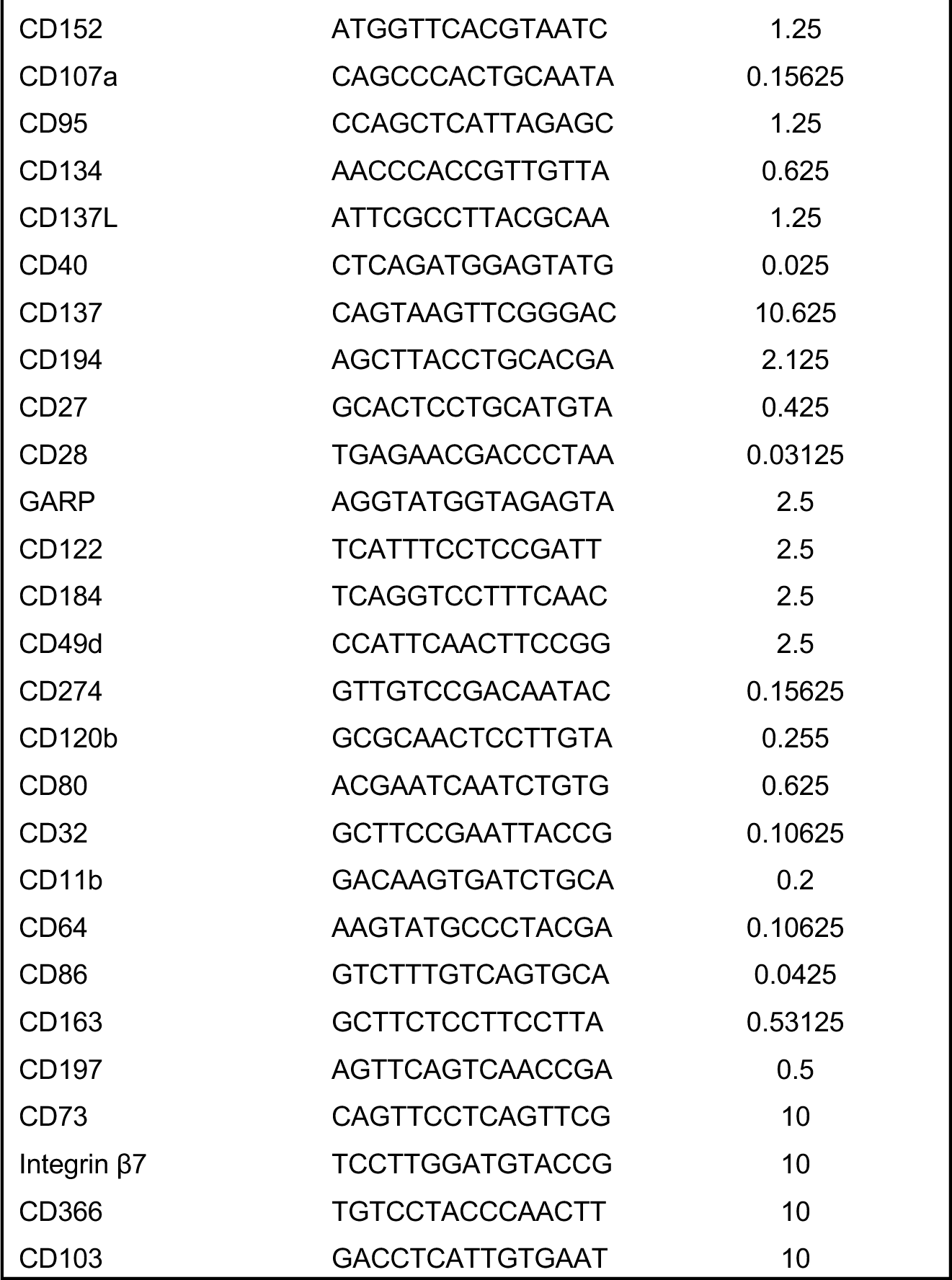
CITE-Seq TotalSeqC antibody details

**Supplemental Table 2:**
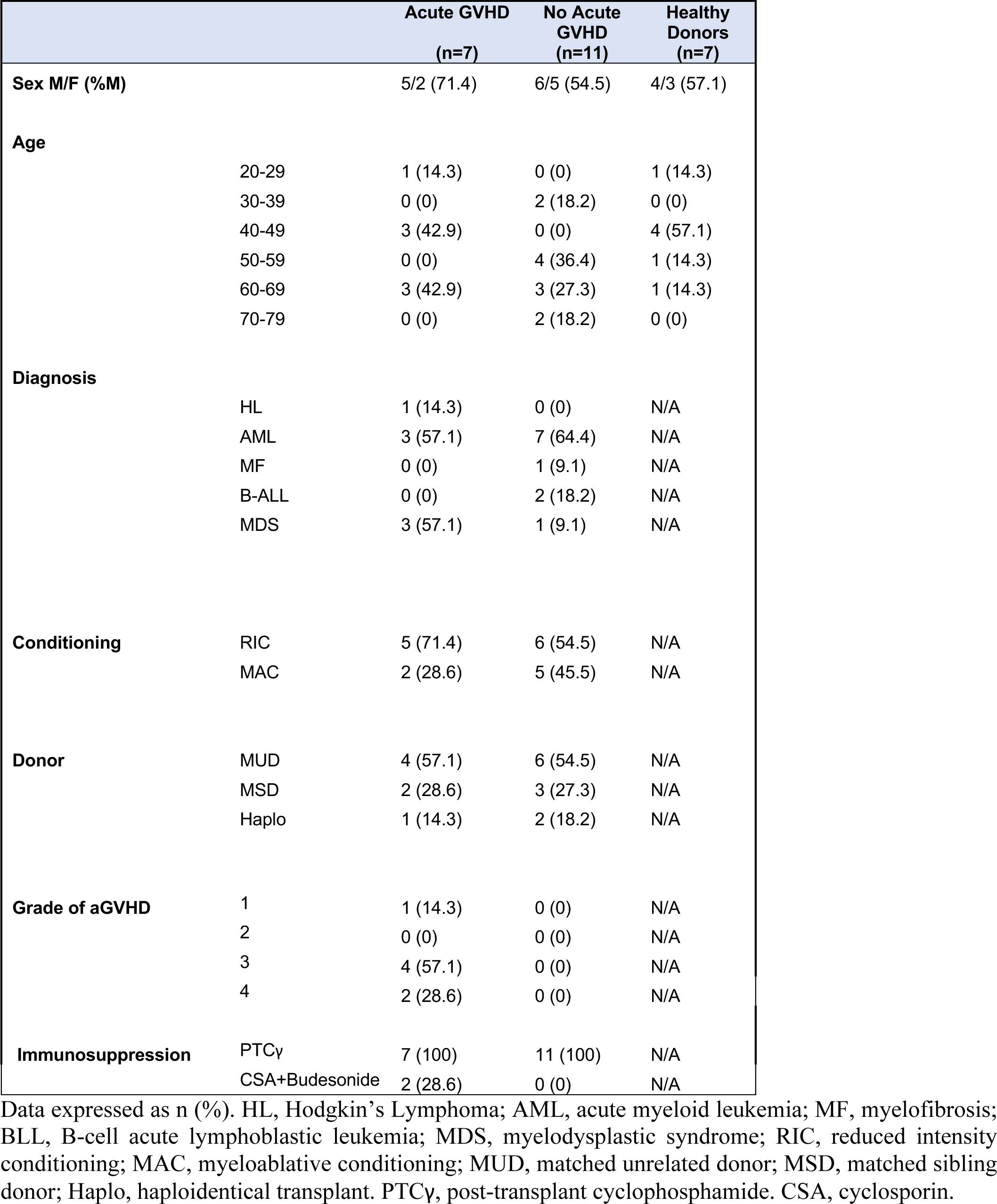
Patient and Healthy Donor Characteristics

**Supplemental Table 3:** Antibodies used for flow cytometry

| Target | Fluorophore | Clone | Company | Dilution |
| --- | --- | --- | --- | --- |
| Amphiregulin | PE | AREG559 | eBioscience | 1:25 |
| CCR6 | APC | G034e3 | BioLegend | 1:25 |
| CCR6 | APC/Fire750 | G034e3 | BioLegend | 1:50 |
| CD117 | PE-Cy7 | 1D11 | BioLegend | 1:50 |
| CD117 | BV650 | 104D2 | BioLegend | 1:50 |
| CD123 | BV510 | 6H6 | BioLegend | 1:100 |
| CD127 | PE | HIL-7R-M21 | BD | 1:50 |
| CD138 | FITC | MI15 | BioLegend | 1:100 |
| CD138 | BV510 | MI15 | BioLegend | 1:100 |
| CD14 | FITC | M5E2 | BioLegend | 1:200 |
| CD14 | BV510 | M5E2 | BioLegend | 1:200 |
| CD15 | FITC | W6D3 | BioLegend | 1:200 |
| CD16 | PE-Dazzle594 | 3G8 | BioLegend | 1:100 |
| CD16 | BUV737 | 3G8 | BD | 1:100 |
| CD19 | FITC | HIB19 | BioLegend | 1:200 |
| CD19 | BV510 | HIB19 | BioLegend | 1:200 |
| CD20 | FITC | 2H7 | BioLegend | 1:200 |
| CD20 | BV510 | 2H7 | BioLegend | 1:200 |
| CD203c | FITC | NP4D6 | BioLegend | 1:100 |
| CD25 | APC | BC86 | eBioscience | 1:25 |
| CD3 | FITC | OKT3 | BioLegend | 1:100 |
| CD3 | FITC | UCHT1 | BioLegend | 1:100 |
| CD3 | BUV395 | UCHT1 | BD | 1:100 |
| CD3 | BUV496 | SK7 | BD | 1:50 |
| CD3 | BV510 | OKT3 | BioLegend | 1:100 |
| CD303 | BV510 | 201A | BioLegend | 1:100 |
| CD33 | FITC | HIM3-4 | BioLegend | 1:100 |
| CD33 | BV510 | P67.6 | BioLegend | 1:100 |
| CD34 | FITC | 581 | BioLegend | 1:100 |
| CD34 | BV510 | 581 | BioLegend | 1:100 |
| CD39 | PerCP-eF710 | eBioA1 (A1) | eBioscience | 1:100 |
| CD4 | FITC | RPA-T4 | BioLegend | 1:200 |
| CD4 | BV605 | RPA-T4 | BD | 1:200 |
| CD45 | BV711 | HI30 | BioLegend | 1:100 |
| CD45 | BUV395 | HI30 | BD | 1:100 |
| CD49d | SuperBright600 | 9F10 | eBioscience | 1:50 |
| CD56 | BV605 | HCD56 | BioLegend | 1:50 |
| CD56 | APC-Cy7 | HCD56 | BioLegend | 1:50 |
| CD73 | BUV737 | AD2 | BD | 1:50 |
| CD79 | FITC | HM47 | BioLegend | 1:100 |
| CD8a | FITC | RPA-T8 | BioLegend | 1:200 |
| CD8a | BV650 | RPA-T8 | BD | 1:50 |
| CD8a | AF700 | RPA-T8 | eBioscience | 1:100 |
| CD86 | BUV737 | 2331 (FUN1) | BD | 1:100 |
| CD94 | PerCP/Cy5.5 | DX22 | BioLegend | 1:100 |
| CRTh2 | BV421 | BM16 | BioLegend | 1:25 |
| CRTh2 | PE-Dazzle594 | BM16 | BioLegend | 1:25 |
| CTLA-4 | PE-Cy7 | 14D3 | eBioscience | 1:50 |
| CXCR3 | BV421 | G025H7 | BioLegend | 1:50 |
| FCεR1a | FITC | AER-37 | BioLegend | 1:100 |
| FCεR1a | BV510 | AER-37 | BioLegend | 1:100 |
| Fixable Viability Dye eF450 | eF450 | -- | eBioscience | 1:1000 |
| Fixable Viability Dye eF506 | eF506 | -- | eBioscience | 1:1000 |
| Fixable Viability Stain 700 | FVS700 | -- | BD | 1:1000 |
| GM-CSF | APC | BVD2-21C11 | BioLegend | 1:50 |
| IFN-γ | BUV395 | B27 | BD | 1:100 |
| IL-10 | AF488 | JES3-9D7 | eBioscience | 1:25 |
| IL-10 | PE | JES3-9D7 | BioLegend | 1:25 |
| IL-13 | BV711 | JES10-5A2 | BD | 1:50 |
| IL-17A | BV786 | N49-653 | BD | 1:25 |
| IL-17F | PE-CF594 | 033-782 | BD | 1:25 |
| IL-22 | eF450 | 22URTI | eBioscience | 1:25 |
| IL-4 | PE-Cy7 | MP4-25D2 | BioLegend | 1:100 |
| IL-9 | PerCP-eF710 | MH9D1 | eBioscience | 1:50 |
| Ki-67 | FITC | 20Raj1 | eBioscience | 1:25 |
| KLRG1 | BV786 | 2F1/KLRG1 | BioLegend | 1:50 |
| NKG2D | PerCP/Cy5.5 | 1D11 | BioLegend | 1:100 |
| NKp44 | BV786 | p44-8 | BD | 1:50 |
| PD-1 | BV785 | EH12.2H7 | BioLegend | 1:100 |
| TCRαβ | FITC | IP26 | BioLegend | 1:200 |
| TCRγδ | FITC | B1 | BioLegend | 1:100 |
| TNF-α | AF700 | MAb11 | eBioscience | 1:50 |
| Zombie NIR Fixable Viability Dye | Zombie NIR | -- | BioLegend | 1:1000 |

**Supplemental Table 4:** Lineage Antibodies for ILC Sort

| Target | Fluorophore | Clone | Company | Dilution |
| --- | --- | --- | --- | --- |
| CD138 | FITC | MI15 | BioLegend | 1:100 |
| CD14 | FITC | M5E2 | BioLegend | 1:200 |
| CD15 | FITC | W6D3 | BioLegend | 1:200 |
| CD19 | FITC | HIB19 | BioLegend | 1:200 |
| CD20 | FITC | 2H7 | BioLegend | 1:200 |
| CD203c | FITC | NP4D6 | BioLegend | 1:100 |
| CD3 | FITC | OKT3 | BioLegend | 1:100 |
| CD3 | FITC | UCHT1 | BioLegend | 1:100 |
| CD33 | FITC | HIM3-4 | BioLegend | 1:100 |
| CD34 | FITC | 581 | BioLegend | 1:100 |
| CD4 | FITC | RPA-T4 | BioLegend | 1:200 |
| CD79 | FITC | HM47 | BioLegend | 1:100 |
| CD8a | FITC | RPA-T8 | BioLegend | 1:200 |
| FCεR1a | FITC | AER-37 | BioLegend | 1:100 |
| TCRαβ | FITC | IP26 | BioLegend | 1:200 |
| TCRγδ | FITC | B1 | BioLegend | 1:100 |

**Supplemental Table 5:** Antibodies for ILC sort

| Target | Fluorophore | Clone | Company | Dilution |
| --- | --- | --- | --- | --- |
| CCR6 | APC | G034e3 | BioLegend | 1:25 |
| CD117 | PE-Cy7 | 1D11 | BioLegend | 1:50 |
| CD127 | PE | HIL-7R-M21 | BD | 1:50 |
| CD16 | PE-Dazzle594 | 3G8 | BioLegend | 1:100 |
| CD45 | BV711 | HI30 | BioLegend | 1:100 |
| CD56 | BV605 | HCD56 | BioLegend | 1:50 |
| CD94 | PerCP/Cy5.5 | DX22 | BioLegend | 1:100 |
| CRTh2 | BV421 | BM16 | BioLegend | 1:25 |
| Fixable Viability Stain 700 | FVS700 | -- | BD | 1:1000 |
| NKG2D | PerCP/Cy5.5 | 1D11 | BioLegend | 1:100 |
| NKp44 | BV786 | p44-8 | BD | 1:50 |

**Supplemental Table 6:** Intracellular Antibodies used for Flow Cytometry

| Target | Fluorophore | Clone | Company | Dilution |
| --- | --- | --- | --- | --- |
| Amphiregulin | PE | AREG559 | eBioscience | 1:25 |
| GM-CSF | APC | BVD2-21C11 | BioLegend | 1:50 |
| IFN-γ | BUV395 | B27 | BD | 1:100 |
| IL-10 | AF488 | JES3-9D7 | eBioscience | 1:25 |
| IL-10 | AF488 | JES3-9D7 | BioLegend | 1:25 |
| IL-10 | PE | JES3-9D7 | BioLegend | 1:50 |
| IL-13 | BV711 | JES10-5A2 | BD | 1:50 |
| IL-17A | BV786 | N49-653 | BD | 1:25 |
| IL-17F | PE-CF594 | 033-782 | BD | 1:25 |
| IL-22 | eF450 | 22URTI | eBioscience | 1:25 |
| IL-4 | PE-Cy7 | MP4-25D2 | BioLegend | 1:100 |
| IL-9 | PerCP-eF710 | MH9D1 | eBioscience | 1:50 |

## List of Abbreviations

ADT: Antibody derived tags
aGVHD: Acute Graft-Versus-Host Disease
AML: Acute myeloid leukemia
BMT: Bone marrow transplant
CBA: Cytometric Bead Array
CITE-Seq: Cellular Indexing of Transcriptomes and Epitopes by Sequencing
EOMES: Eomesodermin
FACS: Fluorescence Activated Cell Sorting
GVL: Graft-Versus-Leukemia Effect
HSCT: Hematopoietic Stem Cell Transplant
IHC: Immunohistochemistry
IL: Interleukin
ILCs: innate lymphoid cells
ILC1: Group 1 innate lymphoid cells
ILC2: Group 2 innate lymphoid cells
ILC2_10_: IL-10-producing ILC2s
ILC3: Group 3 innate lymphoid cells
KIR: Killer Cell Immunoglobulin-like Receptors
LTi: Lymphoid Tissue Inducer Cells
MDSC: Myeloid derived suppressor cell
MSC: Mesenchymal stromal cell
NK Cells: Natural Killer cells
NSG: NOD-scid-IL2Rg^null^
PMA: Phorbol 12-myristate 13-acetate
RT: Room Temperature
S.I.: Small Intestine
sLT: Surface Lymphotoxin
Tc1: Cytotoxic T cell type 1
TGF-β: Transforming Growth Factor Beta
Th1: T helper 1
Th17: T helper 17
Tc1: Type 1 CD8+ cytotoxic T cell
Tr1: Type 1 Regulatory T cell
Treg: Regulatory T cell
XenoGVHD: Xenogeneic Graft-versus-Host Disease

## Acknowledgments

First and foremost, we would like to thank the HSCT patients at UHN who made this work possible. We would also like to thank the nurses and physicians and surgeons at Princess Margaret Cancer Centre for efforts to obtain samples, and acknowledge technical support provided by the Princess Margaret Genomics Centre, particularly Troy Ketela and Julissa Tsao, the Princess Margaret Cancer Centre flow cytometry core and the UHN Pathology Research Program laboratory. We thank Drs Slava Epelman and Clint Robbins for helpful feedback on the manuscript. Schematic figures were created with BioRender.com. KTR is supported by a MITACs Accelerate award. SJC was supported by a Banting and Best Novo Nordisk Scholarship and a Queen Elizabeth II/Dr. Dina Gordon Malkin Graduate Scholarship in Science and Technology. JMM is supported by Ontario Graduate Scholarships, QE II/Aventis Pasteur Graduate Scholarship and a Peterborough. K.M. Hunter Foundation Scholarship. JM is supported by the Gloria and Seymour Epstein Chair in Cell Therapy and Transplantation. SQC is a Tier 2 Canada Research Chair in Tissue-Specific Immune Tolerance. This research was supported by funding from the Canadian Institutes for Health Research (169084) and Natural Sciences and Engineering Research Council of Canada. SQC was also supported by the Medicine by Design program (Canada First Research Excellence Fund), the Ajmera Transplant Centre and Canada Foundation for Innovation grant 38308. We are grateful to Canadian Blood Services and donors for providing research samples for completion of this project. The reporting and interpretation of research findings are the responsibility of the authors, the views expressed herein do not necessarily represent the view of Canadian Blood Services.

## Author contributions

KTR, ASC, JM and SQC designed patient-based studies. KTR, JBL, LZ and SQC designed and implemented xenogeneic GVHD studies. JM established the Hans Messner biobank and the infrastructure required. ASC, INB, TAM and JM liaised with patient care teams to obtain human samples for study. KTR, JAM, JMM, SJC and SQC developed ILC isolation and expansion methods. KTR, SJC, JAM, JMM, DTB, WC, NS, JA, and YX performed experiments and analysed data. KTR, SJC, JMM and SQC designed and implemented the bioinformatic pipeline for scCITEseq. SQC, JBL, AM, LZ and JM supervised the work. KTR and SQC wrote the manuscript, which all authors reviewed and edited.

## Materials and correspondence

Correspondence and requests for materials should be addressed to Sarah Crome.

## References

1. Lim AI et al. Systemic Human ILC Precursors Provide a Substrate for Tissue ILC Differentiation. Cell 2017;168(6):1086–1100.e10.

2. Spits H, Mjösberg J. Heterogeneity of type 2 innate lymphoid cells. Nat Rev Immunol 2022;22(11):701–712.

3. Björkström NK, Kekäläinen E, Mjösberg J. Tissue-specific effector functions of innate lymphoid cells. Immunology 2013;139(4):416–27.

4. Vivier E et al. Innate Lymphoid Cells: 10 Years On. Cell 2018;174(5):1054–1066.

5. Jegatheeswaran S, Mathews JA, Crome SQ. Searching for the Elusive Regulatory Innate Lymphoid Cell. J Immunol 2021;207(8):1949–1957.

6. Rauber S et al. Resolution of inflammation by interleukin-9-producing type 2 innate lymphoid cells. Nat Med 2017;23(8):938–944.

7. Bartemes KR, Kephart GM, Fox SJ, Kita H. Enhanced innate type 2 immune response in peripheral blood from patients with asthma. J Allergy Clin Immunol 2014;134(3):671–678.e4.

8. Falquet M, Ercolano G, Jandus P, Jandus C, Trabanelli S. Healthy and Patient Type 2 Innate Lymphoid Cells are Differently Affected by in vitro Culture Conditions. J Asthma Allergy 2021;14:773– 783.

9. Jegatheeswaran S, Mathews JA, Crome SQ. Searching for the Elusive Regulatory Innate Lymphoid Cell. The Journal of Immunology 2021;207(8):1949–1957.

10. Seehus CR et al. Alternative activation generates IL-10 producing type 2 innate lymphoid cells. Nat Commun 2017;8(1). doi:10.1038/s41467-017-02023-z

11. Howard E et al. IL-10 production by ILC2s requires Blimp-1 and cMaf, modulates cellular metabolism, and ameliorates airway hyperreactivity. Journal of Allergy and Clinical Immunology 2021;147(4):1281–1295.e5.

12. Bando JK et al. ILC2s are the predominant source of intestinal ILC-derived IL-10. Journal of Experimental Medicine 2020;217(2). doi:10.1084/jem.20191520

13. Golebski K et al. Induction of IL-10-producing type 2 innate lymphoid cells by allergen immunotherapy is associated with clinical response. Immunity 2021;54(2):291–307.e7.

14. Derecki NC et al. Meningeal Type-2 Innate Lymphoid Cells Emerge as Novel Regulators of Microglial Activation and Blood-Brain Barrier Stability: A Central Role for IL-10. SSRN Electronic Journal [published online ahead of print: 2019]; doi:10.2139/ssrn.3414004

15. Blanquart E et al. Targeting androgen signaling in ILC2s protects from IL-33–driven lung inflammation, independently of KLRG1. Journal of Allergy and Clinical Immunology 2022;149(1):237–251.e12.

16. Taylor S et al. PD-1 regulates KLRG1+ group 2 innate lymphoid cells. Journal of Experimental Medicine 2017;214(6):1663–1678.

17. Flamar A-L et al. Interleukin-33 Induces the Enzyme Tryptophan Hydroxylase 1 to Promote Inflammatory Group 2 Innate Lymphoid Cell-Mediated Immunity. Immunity 2020;52(4):606–619.e6.

18. Huang Y et al. IL-25-responsive, lineage-negative KLRG1hi cells are multipotential ‘inflammatory’ type 2 innate lymphoid cells. Nat Immunol 2015;16(2):161–169.

19. Garnett C, Apperley JF, Pavlu J. Treatment and management of graft-versus-host disease: Improving response and survival. Ther Adv Hematol 2013;4(6):366–378.

20. Holtan SG et al. Disease progression, treatments, hospitalization, and clinical outcomes in acute GVHD: a multicenter chart review. Bone Marrow Transplant 2022;57(10):1581–1585.

21. Holtan SG et al. Disease progression, hospital readmissions, and clinical outcomes for patients with steroid-refractory acute graft-versus-host disease: A multicenter, retrospective study. Bone Marrow Transplant 2022;57(9):1399–1404.

22. Finazzi MC et al. Characteristics of graft-versus-host disease occurring after alemtuzumab-containing allogeneic stem cell transplants: incidence, organ involvement, risk factors and survival. Br J Haematol 2020;188(4):550–559.

23. Jagasia M et al. Risk factors for acute GVHD and survival after hematopoietic cell transplantation. Blood 2012;119(1):296–307.

24. Luznik L et al. HLA-haploidentical bone marrow transplantation for hematologic malignancies using nonmyeloablative conditioning and high-dose, posttransplantation cyclophosphamide. Biol Blood Marrow Transplant 2008;14(6):641–50.

25. Tang Q, Bluestone JA. Regulatory T-cell therapy in transplantation: moving to the clinic.. Cold Spring Harb Perspect Med 2013;3(11). doi:10.1101/cshperspect.a015552

26. Duggleby R, Danby RD, Madrigal JA, Saudemont A. Clinical Grade Regulatory CD4+ T Cells (Tregs): Moving Toward Cellular-Based Immunomodulatory Therapies. Front Immunol 2018;9:252.

27. Bruce DW et al. Type 2 innate lymphoid cells treat and prevent acute gastrointestinal graft-versus-host disease. Journal of Clinical Investigation 2017;127(5):1813–1825.

28. Piperoglou C et al. Innate lymphoid cell recovery and occurrence of GvHD after hematopoietic stem cell transplantation [Internet]. J Leukoc Biol 2022;111(1):161–172.

29. Kroeze A et al. Presence of innate lymphoid cells in allogeneic hematopoietic grafts correlates with reduced graft-versus-host disease. Cytotherapy 2022;24(3):302–310.

30. Simonetta F, Alvarez M, Negrin RS. Natural killer cells in graft-versus-host-disease after allogeneic hematopoietic cell transplantation. Front Immunol 2017;8(APR). doi:10.3389/fimmu.2017.00465

31. Minculescu L et al. Early Natural Killer Cell Reconstitution Predicts Overall Survival in T Cell– Replete Allogeneic Hematopoietic Stem Cell Transplantation. Biology of Blood and Marrow Transplantation 2016;22(12):2187–2193.

32. Marius Munneke J et al. Activated innate lymphoid cells are associated with a reduced susceptibility to graft-versus-host disease. Blood 2014;124(5):812–821.

33. Huang Q et al. IL -10 producing type 2 innate lymphoid cells prolong islet allograft survival. EMBO Mol Med 2020;12(11). doi:10.15252/emmm.202012305

34. Chen PP et al. Alloantigen-specific type 1 regulatory T cells suppress through CTLA-4 and PD-1 pathways and persist long-term in patients. Sci Transl Med 2021;13(617). doi:10.1126/scitranslmed.abf5264

35. Gagliani N et al. Coexpression of CD49b and LAG-3 identifies human and mouse T regulatory type 1 cells. Nat Med 2013;19(6):739–746.

36. Long A et al. Type 2 Innate Lymphoid Cells Impede IL-33-Mediated Tumor Suppression. J Immunol 2018;201(11):3456–3464.

37. Ohta A et al. A2A Adenosine Receptor May Allow Expansion of T Cells Lacking Effector Functions in Extracellular Adenosine-Rich Microenvironments. The Journal of Immunology 2009;183(9):5487– 5493.

38. Long A et al. Type 2 Innate Lymphoid Cells Impede IL-33–Mediated Tumor Suppression. The Journal of Immunology 2018;201(11):3456–3464.

39. Murphy JM, Ngai L, Mortha A, Crome SQ. Tissue-Dependent Adaptations and Functions of Innate Lymphoid Cells. Front Immunol 2022;13:836999.

40. Mak ML, Reid KT, Crome SQ. Protective and pathogenic functions of innate lymphoid cells in transplantation. Clin Exp Immunol [published online ahead of print: April 29, 2023]; doi:10.1093/cei/uxad050

41. Burman AC et al. IFNγ differentially controls the development of idiopathic pneumonia syndrome and GVHD of the gastrointestinal tract. Blood 2007;110(3):1064–1072.

42. Guy-Grand D, Vassalli P. Gut injury in mouse graft-versus-host reaction. Study of its occurrence and mechanisms. Journal of Clinical Investigation 1986;77(5):1584–1595.

43. Gutiérrez-Hoya A et al. Role of CD8 Regulatory T Cells versus Tc1 and Tc17 Cells in the Development of Human Graft-versus-Host Disease. J Immunol Res 2017;2017. doi:10.1155/2017/1236219

44. Szebeni J, Wang MG, Pearson DA, Szot GL, Sykes M. IL-2 inhibits early increases in serum gamma interferon levels associated with graft-versus-host-disease. Transplantation 1994;58(12):1385–93.

45. Xhaard A et al. Steroid-Refractory Acute GVHD: Lack of Long-Term Improved Survival Using New Generation Anticytokine Treatment. Biology of Blood and Marrow Transplantation 2012;18(3):406–413.

46. Bruce DW et al. Third-party type 2 innate lymphoid cells prevent and treat GI tract GvHD. Blood Adv 2021;5(22):4578–4589.

47. Ehx G et al. Xenogeneic Graft-Versus-Host Disease in Humanized NSG and NSG-HLA-A2/HHD Mice. Front Immunol 2018;9. doi:10.3389/fimmu.2018.01943

48. Proics E et al. Preclinical assessment of antigen-specific chimeric antigen receptor regulatory T cells for use in solid organ transplantation. Gene Ther 2023;30(3–4):309–322.

49. Choi J et al. IFNR signaling mediates alloreactive T-cell trafficking and GVHD [Internet]. 2012:

50. Croudace JE et al. Chemokine-mediated tissue recruitment of CXCR3+ CD4+ T cells plays a major role in the pathogenesis of chronic GVHD. Blood 2012;120(20):4246–55.

51. Crome SQ, Wang AY, Kang CY, Levings MK. The role of retinoic acid-related orphan receptor variant 2 and IL-17 in the development and function of human CD4+ T cells. Eur J Immunol 2009;39(6):1480–1493.

52. Crome SQ et al. Inflammatory Effects of Ex Vivo Human Th17 Cells Are Suppressed by Regulatory T Cells. The Journal of Immunology 2010;185(6):3199–3208.

53. Acosta-Rodriguez E v., et al. Surface phenotype and antigenic specificity of human interleukin 17-producing T helper memory cells. Nat Immunol 2007;8(6):639–646.

54. Cosmi L et al. CRTH2 is the most reliable marker for the detection of circulating human type 2 Th and type 2 T cytotoxic cells in health and disease. Eur J Immunol 2000;30(10):2972–9.

55. Murphy KM, Reiner SL. The lineage decisions of helper T cells. Nat Rev Immunol 2002;2(12):933– 944.

56. Yang L et al. Interleukin-13 interferes with activation-induced t-cell apoptosis by repressing p53 expression. Cell Mol Immunol 2016;13(5):669–77.

57. Reading JL et al. Augmented Expansion of Treg Cells From Healthy and Autoimmune Subjects via Adult Progenitor Cell Co-Culture. Front Immunol 2021;12:716606.

58. Hefazi M, Bolivar-Wagers S, Blazar BR. Regulatory T Cell Therapy of Graft-versus-Host Disease: Advances and Challenges. Int J Mol Sci 2021;22(18). doi:10.3390/ijms22189676

59. McCallion O, Bilici M, Hester J, Issa F. Regulatory T-cell therapy approaches. Clin Exp Immunol 2023;211(2):96–107.

60. Dawson NAJ et al. Systematic testing and specificity mapping of alloantigen-specific chimeric antigen receptors in regulatory T cells. JCI Insight 2019;4(6). doi:10.1172/jci.insight.123672

61. Liu X, Quan N. Immune Cell Isolation from Mouse Femur Bone Marrow. Bio Protoc 2015;

62. Colpitts SJ et al. Strategies for optimizing CITE-seq for human islets and other tissues.. Front Immunol 2023;14:1107582.

63. Hao Y et al. Integrated analysis of multimodal single-cell data. Cell 2021;184(13):3573–3587.e29.

64. McInnes L, Healy J, Saul N, Großberger L. UMAP: Uniform Manifold Approximation and Projection. J Open Source Softw 2018;3(29):861.

65. Blighe K, Rana S, Lewis M. EnhancedVolcano: publication-ready volcano plots with enhanced colouring and labeling. Bioconductor version: Release (3.11), 2020

